# Atypical B cells upregulate co-stimulatory molecules during malaria and secrete antibodies with T follicular helper cell support

**DOI:** 10.1101/2021.12.07.471601

**Authors:** Christine S. Hopp, Jeff Skinner, Sarah L. Anzick, Christopher M. Tipton, Mary E. Peterson, Shanping Li, Safiatou Doumbo, Kassoum Kayentao, Aissata Ongoiba, Craig Martens, Boubacar Traore, Peter D. Crompton

**Affiliations:** Malaria Infection Biology and Immunity Section, Laboratory of Immunogenetics, National Institute of Allergy and Infectious Diseases, National Institutes of Health, Rockville, USA; Rocky Mountain Laboratory Research Technologies Section, Genomics Unit, National Institute of Allergy and Infectious Diseases, National Institutes of Health, Hamilton, USA; Lowance Center for Human Immunology, Division of Rheumatology, Department of Medicine, Emory University School of Medicine, Atlanta, USA; Malaria Research and Training Centre, Department of Epidemiology of Parasitic Diseases, International Center of Excellence in Research, University of Sciences, Techniques and Technologies of Bamako, Mali

## Abstract

Several infectious and autoimmune diseases are associated with an expansion of CD21^-^CD27^-^ atypical B cells (atBCs). The function of atBCs remains unclear and few studies have investigated the biology of pathogen-specific atBCs during acute infection. Here, we performed longitudinal RNA-sequencing and flow cytometry analyses of *Plasmodium falciparum* (*Pf)*-specific B cells before and shortly after febrile malaria, with simultaneous analysis of influenza hemagglutinin (HA)-specific B cells as a comparator. B cell receptor-sequencing showed that *Pf*-specific atBCs, activated B cells (actBCs) and classical memory B cells share clonality and have comparable somatic hypermutation. In response to malaria, *Pf*-specific atBCs and actBCs expanded and upregulated molecules that mediate B-T cell interactions, suggesting that atBCs respond to T follicular helper (Tfh) cells. Indeed, in the presence of Tfh cells and *Staphylococcal enterotoxin B*, atBCs of malaria-exposed individuals differentiated into CD38^+^ antibody-secreting cells *in vitro*, suggesting that atBCs may actively contribute to humoral immunity to infectious pathogens.

**One Sentence Summary:** This study shows that atypical B cells actively respond to acute malaria and have the capacity to produce antibodies with T cell help.

## INTRODUCTION

Infectious diseases such as malaria, HIV-AIDS, TB and SARS-Cov-2-Covid-19, as well as several autoimmune diseases, are associated with an increased frequency of circulating atypical B cells (atBCs) (*1*-*3*). AtBCs express low levels of CD21, lack expression of CD27—the classical human memory B cell (MBC) marker—and upregulate several inhibitory receptors, the integrin CD11c, and express the transcription factor T-bet (*4-8*). Although CD21^-^CD27^-^ B cells differ phenotypically to varying degrees across disease states and have been referred to as ‘atypical’, ‘exhausted’, ‘tissue-like’ or ‘alternative lineage’ B cells, these cells tend to share common features such as the upregulation of inhibitory receptors and diminished B cell receptor (BCR) signaling and cytokine production in response to stimulation *in vitro (5, 9-12*). These findings have led to the hypothesis that atBCs may be anergic or ‘exhausted’ through chronic or repeated exposure to antigen and inflammation.

Malaria remains a serious global health threat, causing an estimated 409,000 deaths and 229 million clinical cases worldwide in 2019 (*13*). More than 90% of malaria cases occur in Africa where *Pf* dominates (*13*). In areas of intense *Pf* transmission, naturally acquired immunity to severe malaria typically develops within the first 5 years of life (*14*), but immunity to uncomplicated febrile malaria is generally not acquired until early adulthood (*14*). Even after decades of *Pf* exposure, individuals rarely appear to develop sterile immunity (*15, 16*). Antibodies are known to play a central role in immunity to *Pf* malaria (*17*), so it has been postulated that the expansion of atBCs may contribute to the inefficient acquisition of humoral immunity to malaria.

Compared to healthy U.S. adults, in whom 2-4% of total circulating B cells are CD21^-^CD27^-^ (*6*), atBCs are increased in individuals residing across diverse malaria-endemic regions (*18-21*). In Mali, for example, atBCs comprise on average 15-20% of total mature B cells in children and up to 40% in healthy adults (*6*). In addition, the degree of atBC expansion appears to correlate with *Pf* transmission intensity (*18, 22*). Exposure of primary human B cells to INF-γ and BCR crosslinking *in vitro* induces the expansion of T-bet^hi^ atBCs (*8*), suggesting that malaria-induced Th1-cytokine responses may contribute to atBC differentiation.

Although one report provided evidence of *Pf*-specific Ig transcripts produced by atBCs *in vivo* and showed that broadly neutralizing *Pf*-specific Abs can be cloned from atBCs (*21*), others have found that atBCs have a reduced capacity to produce antibodies in response to culture conditions that readily induce classical MBCs to secrete antibodies (*5, 12*). In these studies, hyporesponsiveness of atBCs was associated with the expression of inhibitory receptors such as FcγRIIb and Fc receptor-like-5 (FcRL5) (*12*). However, it was shown that *Plasmodium*-specific atBCs proliferate in the mouse infection model (*23*), and that human FcRL5^+^ CD11c^+^ atBCs expand in response to *Pf* infection (*24*). In addition, human atBCs expand in response to *Pf* sporozoite (*Pf*SPZ)-vaccination (*10*) and influenza vaccination (*25*). Finally, FcRL5 is expressed by tetanus vaccine-induced MBCs (*26*), suggesting that atBCs may be related to recently activated BCs (actBCs), although some studies have shown that atBCs and actBC are transcriptionally distinct (*9, 10, 27*).

Here, in a longitudinal cohort in Mali we compared features of *Pf*-specific and influenza hemagglutinin (HA)-specific atBCs, actBCs and MBCs, to gain insight into their function during an acute symptomatic malaria episode. In response to acute malaria, we found that *Pf*-specific atBCs increase in frequency and upregulate molecules that mediate B-T cell interactions. We further found that atBCs of malaria-exposed individuals differentiate into CD38^+^ antibody-secreting cells (ASCs) in the presence of Tfh cells and *Staphylococcal enterotoxin B* (SEB).

## RESULTS

### *Pf*- and HA-specific B cells are not disproportionately enriched for atBCs

Most studies of atBCs in malaria-endemic areas have not examined B cell responses at the antigen specific level. Therefore, the biology of *Pf*-specific atBCs and their frequency relative to atBCs specific for other pathogens remains only partially understood. To address these knowledge gaps we developed flow cytometry–based probes to detect B cells specific for two immunodominant *Pf* proteins: apical membrane antigen 1 (*Pf*AMA1) and merozoite surface protein 1 (*Pf*MSP1), as previously described (*28*). To increase the number of *Pf*-specific B cells detected in each sample, PBMCs were stained simultaneously with *Pf*AMA1 and *Pf*MSP1 probes (Fig. 1A) such that *Pf*AMA1 and *Pf*MSP1 probe-binding cells were indistinguishable and together are referred to as *Pf*^+^ cells. As a comparator non-malaria antigen probe, we used the surface glycoprotein hemagglutinin (HA) of the influenza virus subtypes H1N1 or H3N2, which circulate in Mali (*28, 29*). Influenza appears to circulate year round in West Africa, with two peaks between January–March and August–November (*29*). Having confirmed the specificity of the *Pf* and HA probes (*28*), we first conducted a cross-sectional, *ex vivo* analysis of *Pf*- and HA-specific B cells using cryopreserved PBMCs collected from healthy Malian children (aged 5-17 years; n=34) and adults (aged 18-36 years; n=20) at the end of the 6-month dry season, a period of negligible *Pf* transmission. PBMC samples were obtained from individuals enrolled in the Kalifabougou cohort in Mali where *Pf* transmission is intense and seasonal, as previously described (*15*).

**Figure 1.**
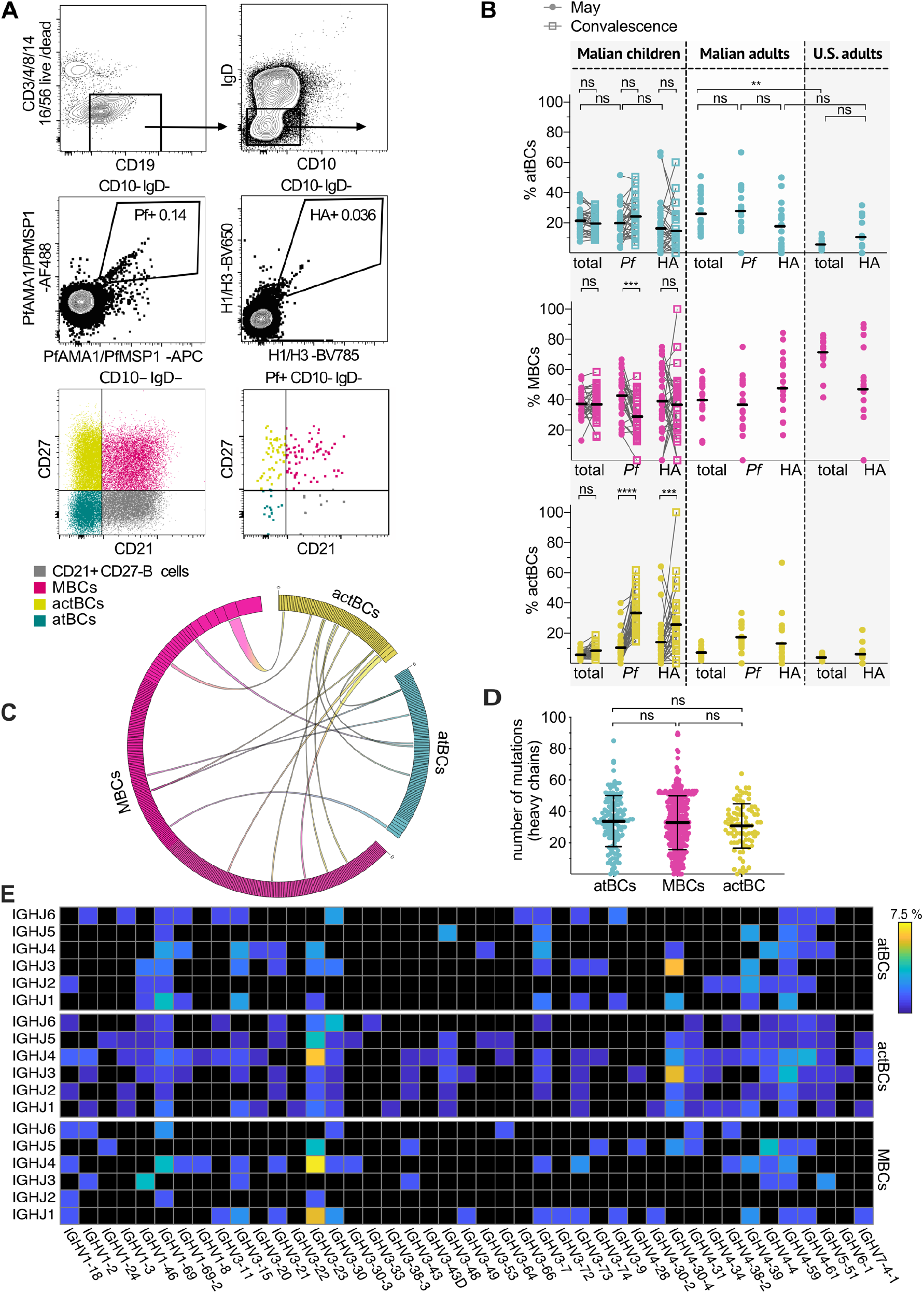
*Pf*- and HA-specific B cells are not disproportionately enriched for atBCs, and *Pf*-specific B cell subsets share clonality. **(A)** Representative flow cytometry plots of antigen-specific B cell subsets. Gating strategy excluded non-B cells (CD3^+^ /CD4^+^/CD8^+^/CD14^+^/CD1^+^/CD56^+^) and IgD^+^ CD10^+^ immature B cells (upper panels). Subsequent gating on *Pf*AMA1 or *Pf*MSP1 (*Pf*^+^) and influenza (HA^+^) probe-binding cells (middle panels). CD21/CD27 staining (bottom panels) within total (left) and *Pf*^+^ (right) CD10-IgD-B cells, with gating to identify actBCs, MBCs and atBCs. **(B)** Percentages of atBCs, MBCs and actBCs within the total, *Pf*^+^ and HA^+^ IgD^-^ CD10^-^ B cell populations in children (5-17 years; n=34) at baseline (filled circles) and 7 days after febrile malaria (empty squares), Malian adults (n=20) and malaria-naïve U.S. adults (n=15) at their baseline. Each dot indicates an individual, connected lines show paired samples, bars show means. Data were acquired from at least five independent experiments. **(C)** Circos plot showing stacked, size-ranked clonal BCR lineages with the largest clones in the clockwise-most position of each population. Connecting lines show matching clones between individual *Pf*^+^ CD10^-^ atBCs (n=143), MBCs (n=371) and actBCs (n=88). **(D)** Number of mutations in the variable region of the heavy chain of single *Pf*^+^ CD10^-^ atBCs, MBCs and actBCs. Data combined from 12 adult Malian donors from at least five independent experiments. Each dot indicates a single cell; lines and whiskers represent means and SDs, respectively. **(E)** Heat map of data displayed in (C), showing usage of Ig heavy chain joining (IGHJ) and variable (IGHV) genes, displayed as a percentage of total sequences obtained from *Pf*^+^ atBCs, MBCs and actBCs. Statistical analysis: (B) Paired comparisons of baseline vs convalescence: 2way ANOVA with Bonferroni’s multiple comparisons test. Remaining comparisons: 1way ANOVA with Tukey’s multiple comparisons test. (D) One-way ANOVA with Tukey’s multiple comparisons test.

We first asked whether *Pf* and HA-specific atBCs exist in this cohort, and if so, whether they differ proportionally from atBCs within the total B cell population. We distinguished B cell subsets based on CD21 and CD27 expression and determined proportions of atBCs (CD21^-^ CD27^-^), MBCs (CD21^+^ CD27^+^) and actBCs (CD21^-^ CD27^+^) within antigen-specific and total CD19^+^ CD10^-^ IgD^-^ mature B cells (Fig. 1A). Consistent with prior studies conducted in Mali (*6, 12, 30*), total atBCs comprised on average 21% of total mature B cells in children and 26% of total mature B cells in adults at the healthy baseline, while only 4% of mature B cells in malaria-naïve U.S. were atBCs (Fig. 1B). The proportion of atBCs within the *Pf*-specific B cell compartment was similar to that of the total B cell population, with an average of 20% in Malian children and 27% in Malian adults. Compared to *Pf*-specific B cells, the average proportion of HA-specific atBCs tended to be lower in Malian children and adults (16% and 17%, respectively), but these differences were not statistically significant (Fig. 1B). On average, 10% of HA-specific B cells were atBCs in U.S. adults, (Fig. 1B), in line with prior reports (*31*). Together these data indicate that *Pf*-specific B cells are not disproportionately atBCs, and that atBCs are not limited to *Pf*-specific B cells in this setting.

### *Pf*- and HA-specific actBCs increase after febrile malaria

Next, in a longitudinal analysis we studied the response of *Pf*- and HA-specific B cells to an acute febrile malaria episode, relative to the healthy baseline. From the same children described above, we analyzed PBMCs collected one week after treatment of their first febrile malaria episode of the 6-month malaria season (convalescence). Although the average percentage of *Pf*- and HA-specific atBCs increased from baseline to convalescence, this was not statistically significant (Fig. 1B). The average percentage of *Pf*-specific actBCs increased from 15% to 30%, while HA-specific actBCs increased to a lesser degree (Fig. 1B). The average percentage of *Pf*-specific MBCs decreased from baseline to convalescence, suggesting that some *Pf*-specific actBCs may originate from resting MBCs in response to acute malaria. B cell subset distributions did not vary significantly with age (Fig. S1).

### *Pf*-specific atBCs, actBCs and MBCs share B cell clones and have comparable somatic hypermutation

To investigate the origin of the *Pf*-specific atBC lineage, we sequenced the BCR of single *Pf*-specific CD19^+^ CD10^-^ atBCs, actBCs and MBCs sorted from PBMCs of healthy Malian adults (n=12) collected at the end of the malaria season. We defined clonality as usage of the same V and J genes, and >85% similarity in a CDR3 region of the same length. We identified clonal relationships between all three B cell subsets (Fig. 1C), suggesting that *Pf*-specific atBCs, actBCs and MBCs are derived from common affinity-matured B cells, in line with what has previously been shown for total non-antigen specific B cell subsets in malaria-exposed individuals (*10*). Next, we estimated heavy chain somatic hypermutation (SHM) frequencies for *Pf*-specific BCRs based on germline BCR sequences in IMGT/V-QUEST (*32*). We found that all three B cell subsets were antigen-experienced and had comparable SHM frequencies (atBCs: 33 ±16; actBCs: 31 ±14; MBCs: 33 ±17 mutations per heavy chain) (Fig. 1D). Analysis of V and J gene usage showed no significant enrichment for a particular V or J gene among *Pf*-specific B cell subsets, although IGHV3-23 and IGHV4-30-4 were frequently used (Fig. 1E).

### *Pf*-specific atBCs are transcriptionally distinct from *Pf*-specific actBCs and MBCs at the healthy baseline

To gain insight into the function of atBCs we performed a transcriptomic analysis of antigen-specific B cell subsets. *Pf*- and HA-specific IgD^-^ CD10^-^ B cells were bulk sorted into atBCs, actBCs and MBCs from PBMCs collected from Malian children (n=16; aged 11-14 years) at their healthy baseline and one week after treatment of their first febrile malaria episode of the ensuing season (see Fig. 2A for sorting strategy). Not all six sorted populations were available for every donor (cell counts ranged from 0-47 cells, see Table S1) and 65 transcriptome libraries were sequenced. Low-expression filters during data processing resulted in the removal of 20 samples, with 45 samples remaining available for paired differential gene expression (DEG) analysis.

**Figure 2.**
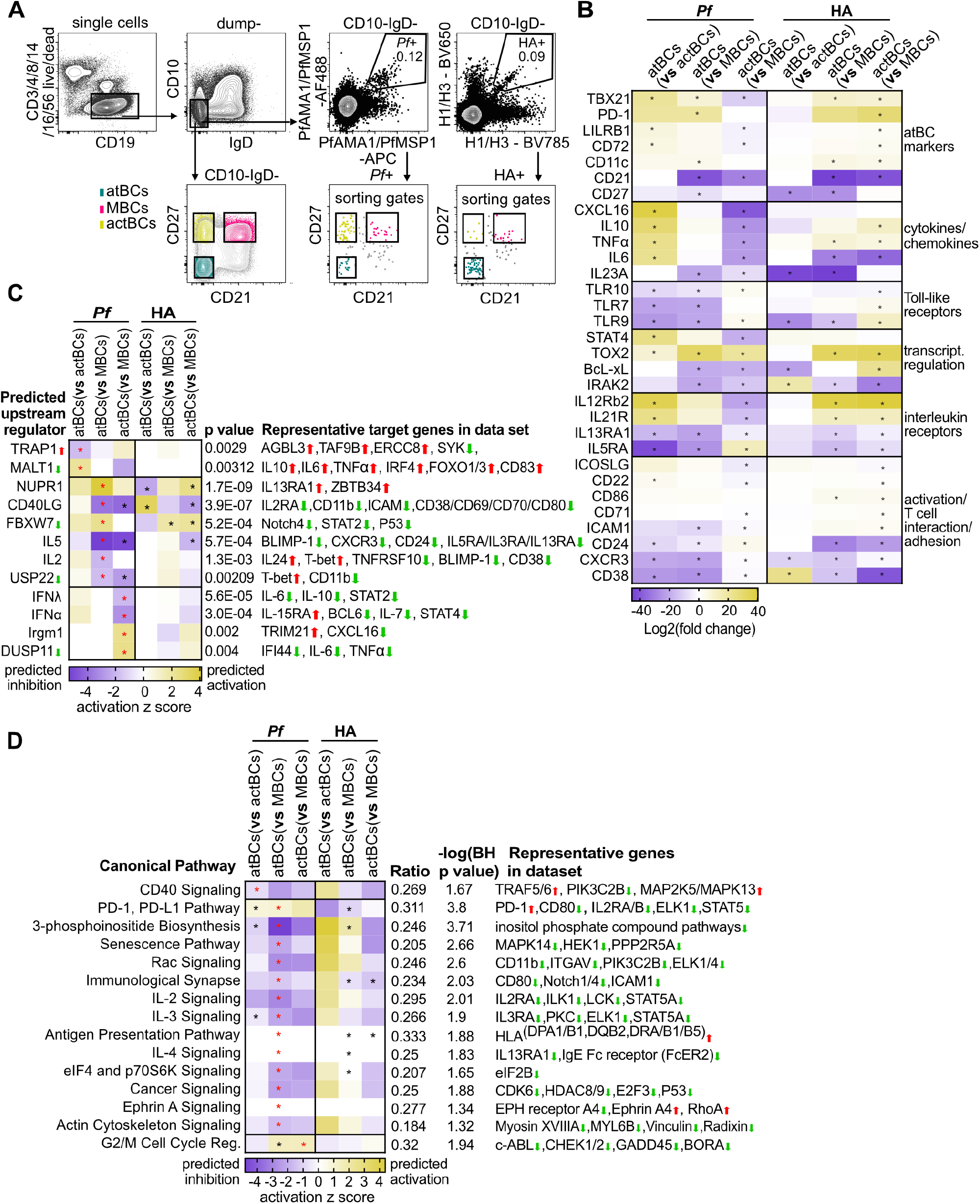
Transcriptomic analysis of *Pf*- and HA-specific B cell subsets at the healthy baseline reveals distinguishing features of *Pf*-specific atBCs. **(A)** Representative flow cytometry plots showing the gating strategy to sort mature CD10-IgD-*Pf*- and HA-specific atBCs, MBCs and actBCs. **(B)** Heatmap showing selected DEGs for the three comparisons (atBCs vs actBCs; atBCs vs MBCs; actBCs vs MBCs) of *Pf*- and HA-specific B cells. Significant DEGs (false discovery rate (FDR)<0.05, no FC cutoff) are indicated with an asterisk. **(C)** Ingenuity upstream regulator analysis using DEGs with FDR<0.2 and no FC cutoff. Heatmap shows predicted regulators with a |z score|>2 and p<0.005 for the respective subset comparison with positive z-score indicating activation of the regulator. Arrow up/down indicates gene expression in the data set. P values <0.005 are indicated with an asterisk; p values and representative target genes pertain to comparisons indicated by red asterisks. **(D)** Canonical pathways that are significantly overlapping with DEGs of the respective subset comparison (FDR<0.2, no FC cut-off) are indicated with asterisks. Ratio indicating the proportion of genes in the canonical pathway that are differentially expressed, the BH-adjusted p value, along with representative genes in the pathway (arrow up/down indicating gene expression) pertain to the comparison indicated with red asterisks. Antigen Presentation pathway, IL-4 signaling, Ephrin A Signaling: no z score can be calculated for these Ingenuity canonical pathways.

We first compared the transcriptional profiles of *Pf*-specific atBCs, actBCs and MBCs at the healthy baseline. *Pf*-specific atBCs expressed higher levels of several genes that have previously been shown to be upregulated by total atBCs (*10, 12*), including *TBX21* (T-bet), *LILRB1, ITGAX* (CD11c) and *CD72*, while CD21 and CD27 were downregulated as expected (Fig. 2B). *Pf*-specific atBCs also expressed higher levels of *PDCD1* (PD-1), and lower levels of *ICAM1, CD24, CD38, CXCR3, TLR7, TLR9* and *TLR10*, relative to *Pf*-specific actBCs and MBCs (Fig. 2B). The transcriptional regulators *STAT4* and *TOX2* were upregulated, while *BcL-xL* and *IRAK2* were downregulated in *Pf*-specific atBCs relative to *Pf*-specific actBCs and MBCs (Fig 2B). *Pf*-specific atBCs also upregulated *CXCL16, IL-10, TNFα* and *IL-6* relative to *Pf*-specific actBCs and displayed an interleukin receptor profile that was distinct from *Pf*-specific actBCs and MBCs, characterized by higher expression of genes encoding the IL-12 and IL-21 receptors and reduced expression of genes encoding the IL-5 and IL-13 receptors (Fig. 2B). Although some features of *Pf*-specific atBCs, such as the downregulation of TLR7 and 10, were unique to *Pf*-specific atBCs, other DEGs were shared by *Pf*-specific and HA-specific atBCs at the healthy baseline.

### Biological pathway and upstream regulator analysis distinguish *Pf*-specific atBCs

We performed an Ingenuity upstream regulators analysis based on DEGs of the following comparisons: atBCs vs actBCs, atBCs vs MBCs, and actBCs vs MBCs. We found that the upregulation of *IL-10, IL-6, TNFα* and *IRF4*, observed in *Pf*-specific, but not HA-specific atBCs, was predicted to be causally linked to the pathway downstream of MALT1 (Fig. 2C). Furthermore, the downregulation of pathways downstream of IL-2 were linked to the upregulation of T-bet uniquely in *Pf*-specific atBCs (Fig. 2C). Next, to identify biological pathways that characterize *Pf*-specific atBCs, actBCs and MBCs, we determined significant overlap between DEGs of the three cell subset comparisons and Ingenuity canonical pathways (Fig. 2D). Compared to *Pf*-specific actBCs and MBCs, *Pf*-specific atBCs exhibit an intracellular signaling profile that suggests an attenuated response to CD40, IL-2, IL-3, IL-4, the B-T cell synapse and actin cytoskeleton signaling (Fig. 2D). In addition, several intracellular signaling molecules/pathways were downregulated in *Pf*-specific, but not HA-specific atBCs, such as the Rac signaling pathway, *ELK1/4, PIK3C2B, MAPK*inases and *STAT5A*, as well as enzymes involved in the generation of inositol phosphate compounds (Fig. 2C), consistent with previous data showing that total atBCs are hyporesponsive to signals that typically activate MBCs (*12*). Finally, the transcriptome analysis suggested that both *Pf*- and HA-specific atBCs have an increased capacity for antigen presentation through upregulation of *HLA-DPA1, -B1, -DQB2, -DRA, -B1*, and *-B5* (Fig. 2D).

### *Pf*-specific atBCs and actBCs have a similar transcriptional response to acute malaria

Next, we sought to gain insight into how *Pf*-specific atBCs respond functionally to acute malaria by comparing antigen-specific B cell subset transcriptomes one week after treatment of acute febrile malaria to the baseline transcriptomes described above. Paired samples from the same 16 children described above were used for this analysis. Transcriptome libraries (n=91) were sequenced and data from 16 samples were removed by low expression filters in the subsequent data processing, leaving 75 samples available for paired analysis of DEGs using DESeq2. Relative to baseline, febrile malaria induced the greatest number of transcriptional changes in *Pf-*specific atBCs, while *Pf-*specific atBCs and actBCs had the greatest overlap in DEGs (Fig. 3A). *Pf-*specific MBCs appeared comparatively quiescent, possibly because activation of resting MBCs downregulates CD21 (*33*), thus excluding responsive MBCs from the MBC gate during the flow sort. Unsupervised hierarchical clustering showed that *Pf-*specific atBCs and actBCs were transcriptionally similar to each other and distinct from *Pf-*specific MBCs, particularly in response to acute malaria (Fig. 3B). This trend was seen for both the *Pf*- and HA-specific B cell subsets (Fig. 3B).

**Figure 3.**
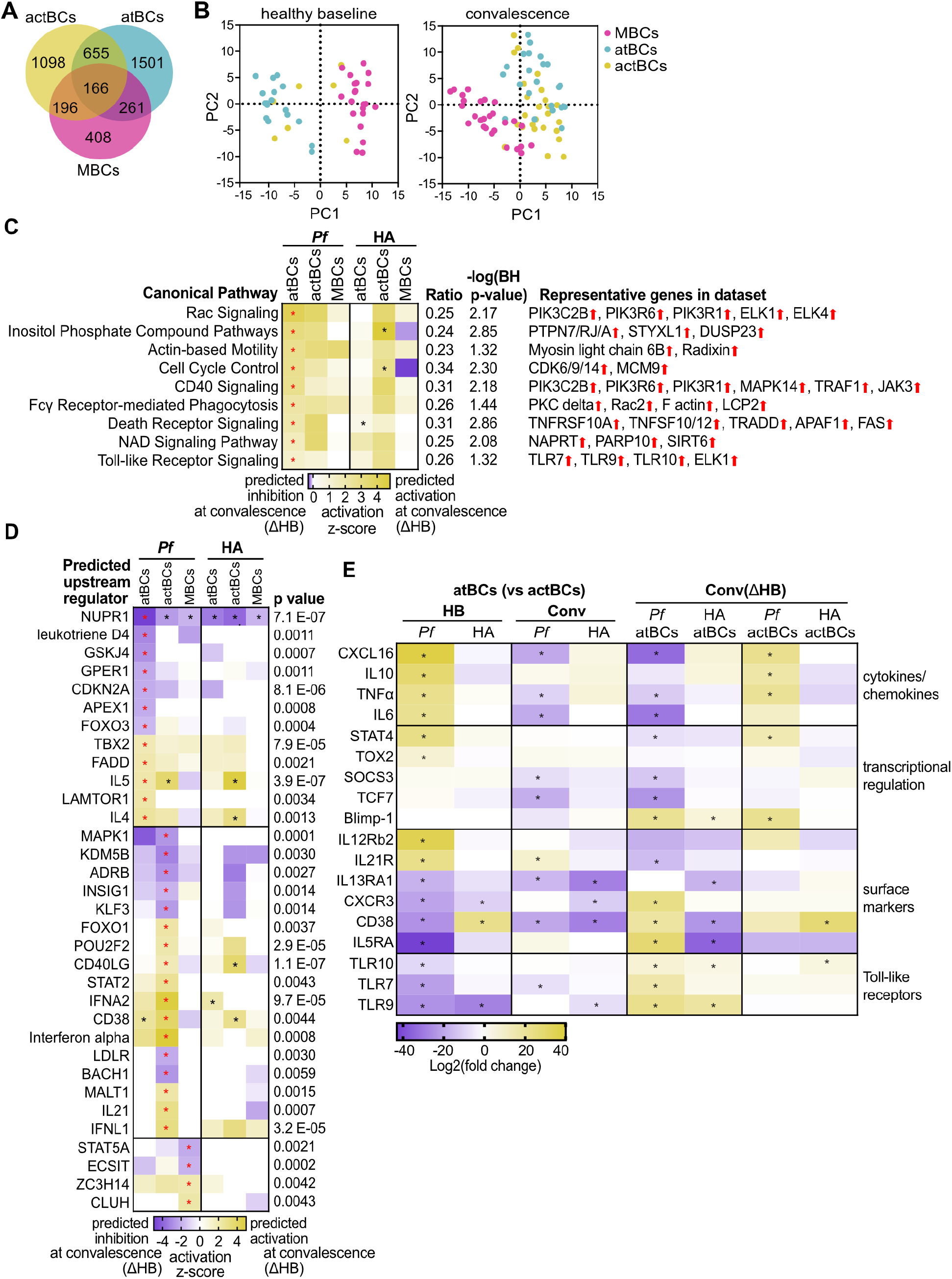
Pathways and upstream regulators defining the response of *Pf*-specific atBCs to malaria are frequently shared by *Pf*-specific actBCs. **(A)** Venn diagram showing the number of DEGs uniquely regulated in response to malaria (Δ healthy baseline, HB) in *Pf*-specific subsets (cut-off FDR<0.05; |FC|>2). **(B)** Principal component analysis using the top 250 most variably expressed genes among *Pf*-specific B cell subsets at baseline and one week after malaria treatment. Each dot represents one antigen-specific B cell subset per donor; *Pf*- and HA-specific samples were pooled to increase the number of data points per plot. **(C)** Canonical pathways that are significantly overlapping with DEGs of *Pf*-specific atBCs at convalescence (Δ HB; FDR<0.2, no FC cut-off). Ratio of overlap, BH-adjusted p value, as well as representative genes in the data set (arrow up/down indicating gene expression) pertain to the comparison indicated with red asterisks. No significant canonical pathway enrichment was found for *Pf*-specific actBCs and MBCs at convalescence (Δ baseline). **(D)** Top upstream regulators identified by Ingenuity analysis using the DEGs of B cell subsets at convalescence (Δ HB; FDR<0.2, no FC cut-off). Heatmap shows predicted regulators with a |z score|>2 and p<0.005 for the respective subset comparison with positive z-score indicating activation of the regulator. Listed p values pertain to comparison indicated by red asterisks. Arrow up/down indicates gene expression in the data set. **(E)** Heatmap showing selected DEGs for the atBCs vs actBCs comparison at HB and convalescence (Conv) on the left and differential expression of these genes in atBCs and actBCs at convalescence (ΔHB) on the right. Significant DEGs (FDR<0.05, no FC cutoff) are indicated with an asterisk.

Of note, several canonical pathways underrepresented in *Pf*-specific atBCs at baseline (Fig. 2C), were overrepresented in response to acute malaria, including Rac-, CD40- and actin cytoskeleton signaling, as well as pathways involving the generation of inositol phosphate compounds (Fig. 3C). In contrast, HA-specific atBCs appeared relatively quiescent in response to malaria (Fig. 3C). We did not identify statistically significant activation or inhibition of pathways among *Pf*-specific actBCs and MBCs in response to acute malaria, although all pathways activated in *Pf*-specific atBCs also trended toward activation in actBCs (Fig. 3C), underscoring the relatedness of these B cell subsets.

Next, we performed an Ingenuity upstream regulator analysis using DEGs of atBCs, actBCs and MBCs at convalescence relative to healthy baseline, to identify regulators that may be involved in orchestrating the response of atBCs to acute malaria. We identified IL-4 and IL-5 as potential upstream activators of atBCs following acute malaria (Fig. 3D). Transcriptional regulators such as *NUPR1* and *FOXO3* were predicted to be negative regulators of atBCs in response to malaria while *TBX2* was a predicted positive regulator (Fig. 3D), consistent with earlier studies of total atBCs (*8, 12*). Although not consistently statistically significant, several of these upstream regulators trended in the same direction in actBCs (Fig. 3D). Except for NUPR1, these predicted upstream regulators were unique to *Pf*-specific atBCs and not observed in HA-specific atBCs (Fig. 3D). In summary, this analysis reveals that *Pf*-specific atBCs and actBCs share a similar transcriptional response to acute malaria in children. This is in line with previous observations that malaria-associated total atBCs have an activated phenotype that manifests as elevated levels of phosphorylated kinases (*12*).

### *Pf*-specific atBCs upregulate *IL-5RA, CD38, PRDM1, CXCR3, TLR7, 9 and 10* in response to acute malaria

While many DEGs were shared by *Pf*-specific atBCs and actBCs, certain DEGs that distinguished *Pf*-specific atBCs at baseline were uniquely regulated in *Pf*-specific atBCs in response to malaria. For example, *CXCL16, TNFα, IL-6, STAT4* and *IL21R* were expressed at higher levels in *Pf*-specific atBCs relative to actBCs at baseline, and then were uniquely downregulated by *Pf*-specific, but not HA-specific atBCs in response to malaria (Fig. 3E). Conversely, *IL-5RA, CD38, CXCR3, TLR7, 9* and *10* were expressed at lower levels in *Pf*-specific atBCs compared to actBCs at baseline, and then were upregulated by *Pf*-specific atBCs in response to malaria. Except for TLR9, these trends were unique to *Pf*-specific atBCs. *PRDM1*, which encodes for B-lymphocyte-induced maturation protein 1 (BLIMP1), and *CD38* were upregulated in *Pf*-specific atBCs and actBCs in response to malaria (Fig. 3E), suggesting activation of pathways involved in differentiation into ASCs.

### *Pf*-specific atBCs and actBCs express markers indicative of T cell interactions

Previous studies found that total atBCs upregulate proteins involved in antigen presentation to T cells, including ICOSL, CD86 and HLA-DR (*9, 12, 34*). To determine whether malaria upregulates the expression of these proteins on *Pf*-specific atBCs, we performed flow cytometry analysis on paired PBMC samples collected from 2–17-year-olds at baseline before the malaria season, and one week after treatment of their first malaria episode of the ensuing malaria season. At baseline both total and *Pf*-specific atBCs expressed higher levels of ICOSL compared to *Pf*-specific actBCs and MBCs (Fig. 4A), in line with the transcriptomic data (Fig. 4H). However, in response to malaria ICOSL was downregulated at the protein and RNA levels in *Pf*-specific atBCs (Fig. 4A&H, Conv(HB)). Both total and *Pf*-specific atBCs and actBCs upregulated CD86 in response to malaria relative to baseline (Fig. 4B). RNA-seq analysis also showed that *Pf*-specific atBCs upregulate *HLA-DPA1, -B1, -DQB2, -DRA, - B1* and *-B5* relative to MBCs (Fig. 2D). By flow cytometry we confirmed that HLA-DR was expressed at significantly higher levels on *Pf*-specific atBCs and actBCs compared to *Pf*-specific MBCs in response to malaria (Fig. 4C). Collectively, these data suggest that *Pf*-specific atBCs upregulate molecules that mediate interactions with CD4 T cells. Moreover, the percentage of *Pf*-specific atBCs that were Ki67^+^ increased significantly at convalescence (Fig. 4D), which taken together with the upregulation of CXCR3 and the lack of significant expansion of peripheral *Pf*-specific atBCs at convalescence (Fig. 1B), suggests that *Pf*-specific atBCs may be recruited to secondary lymphoid organs.

**Figure 4:**
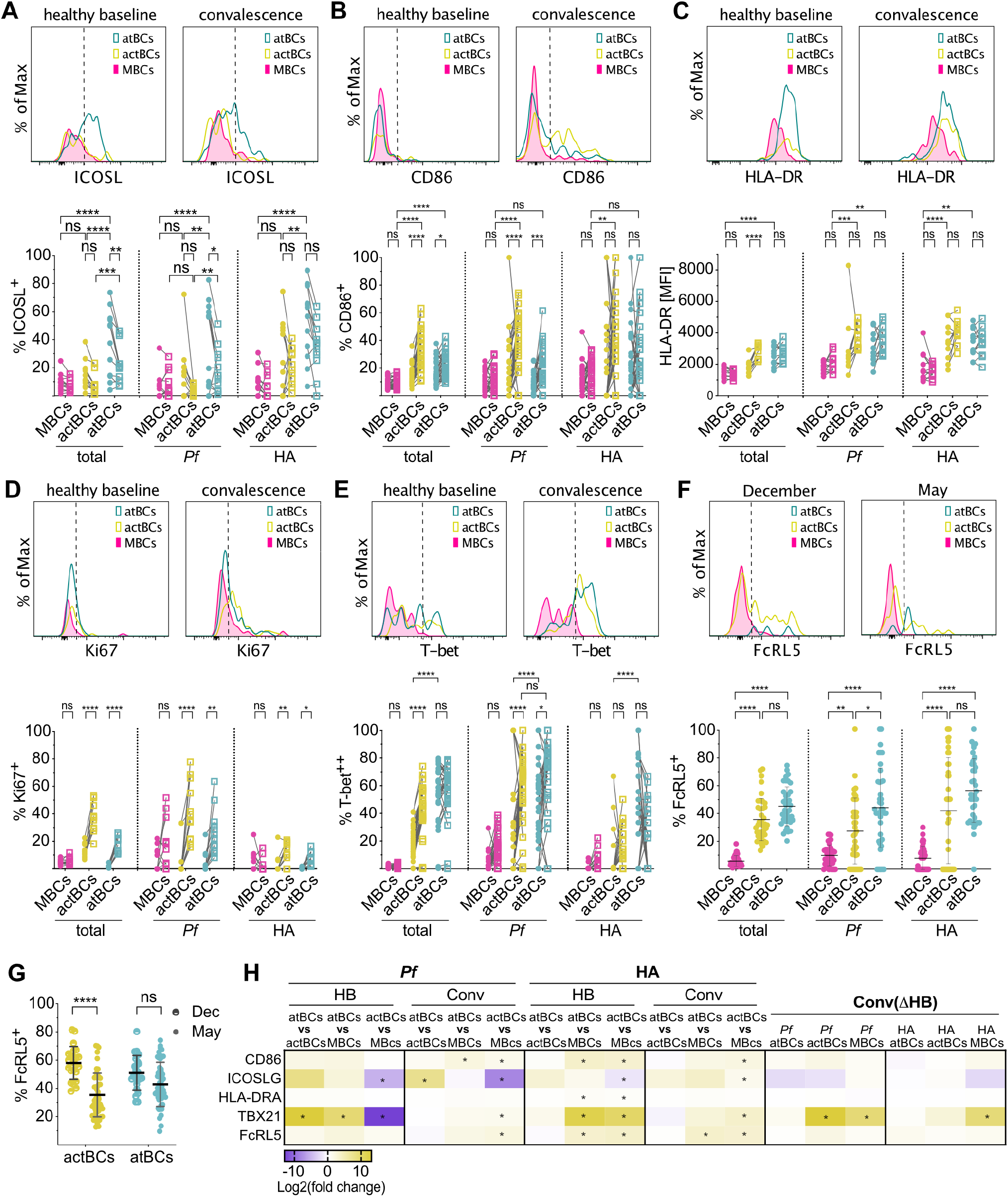
*Pf*-specific atBCs and actBCs express markers indicative of proliferation and T cell interaction. (**A-E**) Histograms (top panels) showing representative mean fluorescence intensity (MFI) values of ICOSL, CD86, HLA-DR, Ki67 and T-bet for *Pf*-specific CD10-IgD-B cell subsets at baseline and one week after treatment of febrile malaria (convalescence). Graphs (bottom panels) show either percentages of cells expressing ICOSL (A), CD86 (B), Ki67 (D) and T-bet (E), or MFIs of HLA-DR (C) for total, *Pf*- and HA-specific CD10-IgD-MBCs (pink), actBCs (yellow) and atBCs (teal) at baseline (filled circles) and convalescence (empty squares). Ki67, ICOSL and HLA-DR data: donors 11-17 years (n=12); CD86 data: donors 2-17 years (n=33). T-bet data: donors 5-17 years (n=33). **(F)** Histograms (top panels) showing representative MFI values of FcRL5 for *Pf*-specific CD10-IgD-B cell subsets at the end of the 6-month malaria season (December) and at baseline (May). Graphs (bottom panels) show percentages of cells expressing FcRL5 for total, *Pf*- and HA-specific CD10-IgD-MBCs (pink), actBCs (yellow) and atBCs (teal) in December (filled circles) and May (empty squares). Donors 13-15 years (n=39). **(G)** Frequency of total atBCs or actBCs expressing FcRL5 in December and after five months of dry season in May. Each dot indicates an individual, lines represent means. Donors 13-15 years (December n=23; May n=39). **(H)** Heatmap showing RNA expression of selected genes. Differential expression analysis and statistical analysis of comparisons were performed using DESeq2. Significant DEGs (FDR<0.05, no FC cutoff) are indicated with an asterisk. Statistical analysis: (A-E) two-way ANOVA with Tukey’s multiple comparisons test. (F-G) one-way ANOVA with Holm-Sidak’s multiple comparisons test.

### *Pf*-specific atBCs and actBCs upregulate T-bet and FcRL5 after malaria, while only atBCs maintain expression of these markers through the dry season

Previous studies have shown that T-bet is upregulated in total atBCs of malaria-exposed children and adults (*8*). Here, we found by flow cytometry that T-bet expression is upregulated at baseline in both *Pf*-specific atBCs and actBCs relative to classical MBCs, although a significantly higher proportion of total, *Pf*- and HA-specific atBCs were T-bet^hi^ compared to actBCs (Fig. 4E). In response to acute malaria the proportion of T-bet^hi^ cells increased for both *Pf*-specific atBCs and actBCs, with the increase being particularly marked for *Pf*-specific actBCs (Fig. 4E). These results approximate the RNA-seq data that showed higher *TBX21* expression in *Pf* -specific atBCs compared to *Pf*-specific actBCs and MBCs at baseline, and upregulation of *TBX21* in *Pf* -specific actBCs and MBCs in response to malaria (Fig. 4H). However, in contrast to *Pf* -specific actBCs, *Pf* -specific atBCs appear to maintain high T-bet expression in the absence of ongoing exposure to malaria (i.e. at baseline at the end of the 6-month dry season) (Fig. 4E&H).

Similarly, FcRL5, another marker of atBCs, was upregulated at baseline in both *Pf*-specific atBCs and actBCs relative to classical MBCs, although a significantly higher proportion of *Pf*-specific atBCs were FcRL5^+^ compared to *Pf*-specific actBCs (Fig. 4F). To assess the stability of FcRL5 expression in the absence of ongoing malaria exposure, we analyzed B cells collected from the same individuals at the beginning (December) and end (May) of the dry season when *Pf* transmission is negligible. While the proportion of FcRL5^+^ *Pf* -specific atBCs and actBCs both decreased over the dry season, the decrease was only statistically significant for *Pf* -specific actBCs (Fig. 4G).

In summary, *Pf*-specific atBCs appear more likely to retain T-bet and FcRL5 expression even at homeostasis, in contrast to *Pf* -specific actBCs that readily upregulate these molecules in response to malaria, but then partially lose their expression in the absence of ongoing malaria exposure.

### AtBCs upregulate CD38 and secrete IgG and IgM in the presence of Tfh cells

In the germinal center (GC), Tfh cells provide critical support for the differentiation of naive B cells into isotype-switched, affinity-matured long-lived plasma cells (LLPCs) and classical MBCs through contact-dependent and soluble signals including the T cell receptor, ICOS, CD28, CD40L and IL-21 (*35*). Whether Tfh cells can drive atBCs to differentiate into ASCs is unknown. Based on our observation that *Pf*-specific atBCs express higher levels of HLA-DR and ICOSL relative to MBCs and upregulate CD86, CD40-signaling and the IL-21 receptor in response to malaria, we hypothesized that atBCs may be poised for productive interactions with Tfh cells. To test this hypothesis we FACS-sorted circulating CD4^+^ CD3^+^ CD45RO^+^ CXCR5^+^ PD-1^+^ Tfh cells (cTfh) (Fig. 5A) that are known to resemble GC-Tfh cells phenotypically and functionally (*36, 37*). We then co-cultured these cTfh cells with autologous atBCs, MBCs or actBCs (Fig. 5B) for 7 days with or without the superantigen *Staphylococcal enterotoxin B* (SEB) to mimic the interaction between antigen-specific T cells and antigen-presenting B cells(*36, 37*). On average, we found that cTfh cells induced expression of the plasmablast marker CD38 in approximately 60% of atBCs, and 90% of MBCs (Fig. 5C-D). Accordingly, cTfh cells induced atBCs to secrete IgM and IgG1-4 (Fig. 5E), albeit at lower levels than that produced by MBCs (Fig. S2).

**Figure 5:**
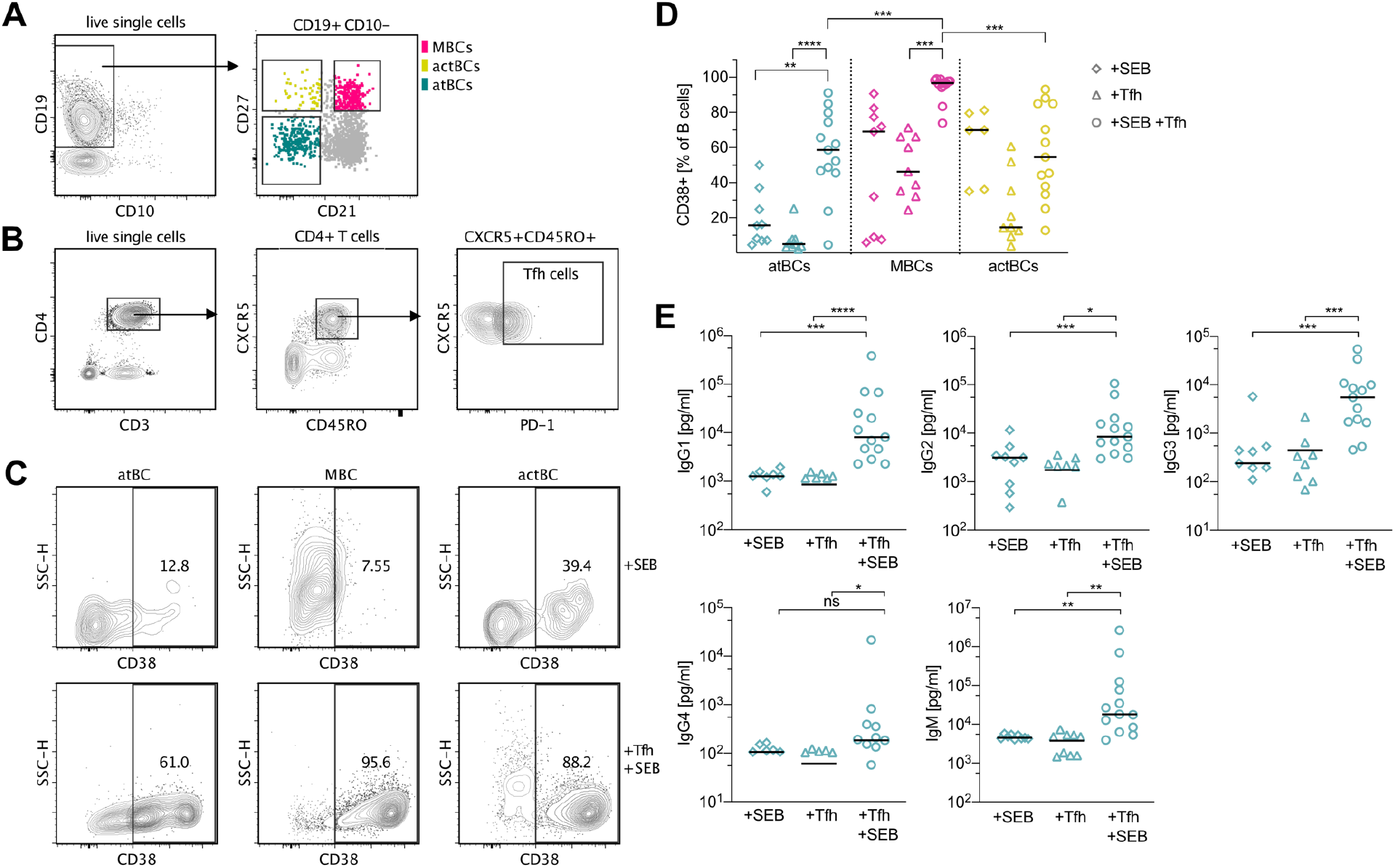
atBCs upregulate CD38 and secrete Ig when co-cultured with PD-1^+^ CXCR5^+^ CD45RO^+^ CD4^+^ cTfh cells. Representative flow cytometry plots showing the gating strategy used to FACS-sort (**A**) mature CD10-B cell subsets as well as (**B**) CD4^+^ CD3^+^ CD45RO^+^ CXCR5^+^ PD-1^+^ cTfh cells. B cell subsets were cultured with or without autologous cTfh cells in the presence or absence of SEB. After 7 days, surface expression of CD38 and secreted Ig levels were measured. (**C**) Representative plots of CD38 staining on B cell subsets after 7-days of co-culture with SEB +/-cTfh cells. (**D**) Percentage of B cell subsets that are CD38^+^ after 7-day co-culture with SEB, cTfh cells, or SEB plus cTfh cells. Symbols represent individual donors (n=13), lines represent medians. (**E**) IgG1–4 and IgM concentration in supernatants after 7-day co-culture with SEB, cTfh cells, or SEB plus cTfh cells. Symbols represent individual donors (n=13), lines represent medians. Statistical analysis: (D-E) Kruskall-Wallis with Dunn’s multiple comparison test. Data are representative of four independent experiments.

## Discussion

atBCs are associated with several infectious and autoimmune diseases, yet their origin and function remain only partially understood, particularly at the level of antigen-specific atBCs. Moreover, in the case of infection-associated atBCs, the extent to which atBCs are specific for the infecting pathogen remains unclear. Here we conducted a longitudinal study of Malian children and adults to investigate the biology of *Pf*- and influenza (HA)-specific atBCs at the homeostatic baseline before the malaria season and in response to an acute symptomatic *Pf* infection during the ensuing malaria season.

Regarding the specificity of atBCs in the context of malaria exposure, we found that the proportions of *Pf-* and HA-specific atBCs were similar, and comparable to the proportion of atBCs within total B cells, indicating that atBCs are not disproportionately *Pf*-specific and are not limited to *Pf*-specific B cells in this setting. In contrast, a recent study in Kenya showed that *Pf-*specific B cells were more likely than tetanus-toxoid (TT)-specific B cells to be atBCs (*22*). The discordant findings between the Kenya study and the present work may be due in part to differences in the frequency of HA exposure (annual) versus TT exposure (childhood vaccine). Aye et al. (*22*) also found that malaria exposure correlated with a higher proportion of *Pf*-specific and TT-specific atBCs, suggesting that malaria drives the atBC phenotype in both *Pf*-specific and bystander B cells (*22*). In malaria-endemic areas it will be of interest to study atBCs specific for a broader range of non-malaria pathogens and vaccine antigens to better define the relationship between the quality and frequency of antigen exposure and atBC differentiation.

Concerning the origin and clonality of atBCs, BCR-seq analysis provided evidence of comparable SHM rates and clonal relationships between *Pf-*specific atBCs, actBCs and MBCs, suggesting that these subsets originate from common affinity-matured B cell precursors. This is consistent with prior studies of bulk B cell subsets in malaria-exposed individuals in Mali (*12*), Kenya (*10*) and Uganda (*38*), but at odds with a study in Gabon that showed little evidence of clonal relationship between atBCs and MBCs (*21*).

RNA-seq analysis of *Pf*-specific atBCs shed light on their phenotype and potential origin and function. *Pf*-specific atBCs have a muted intracellular signaling response to cytokines and T cell interactions, which is consistent with previous functional studies of bulk atBCs (*12*). In addition, *Pf*-specific atBCs at the healthy baseline, relative to MBCs, express higher levels of *STAT4* and the genes encoding the IL-12- and IL-21-receptors. IL-12 has been shown to induce a cascade of events in B cells that is similar to Th1-commitment, including STAT4 activation and upregulation of T-bet- and the IL-12-receptor (*39*). Furthermore, IL-21 drives naïve B cells to a CD27^low^T-bet^+^ FcRL5^+^ IL-21R^+^ CD11c^+^ phenotype (*40*). Together, these findings point toward potential roles for IL-12 and IL-21 in the differentiation of *Pf*-specific atBCs (see model in Fig. 6).

**Figure 6:**
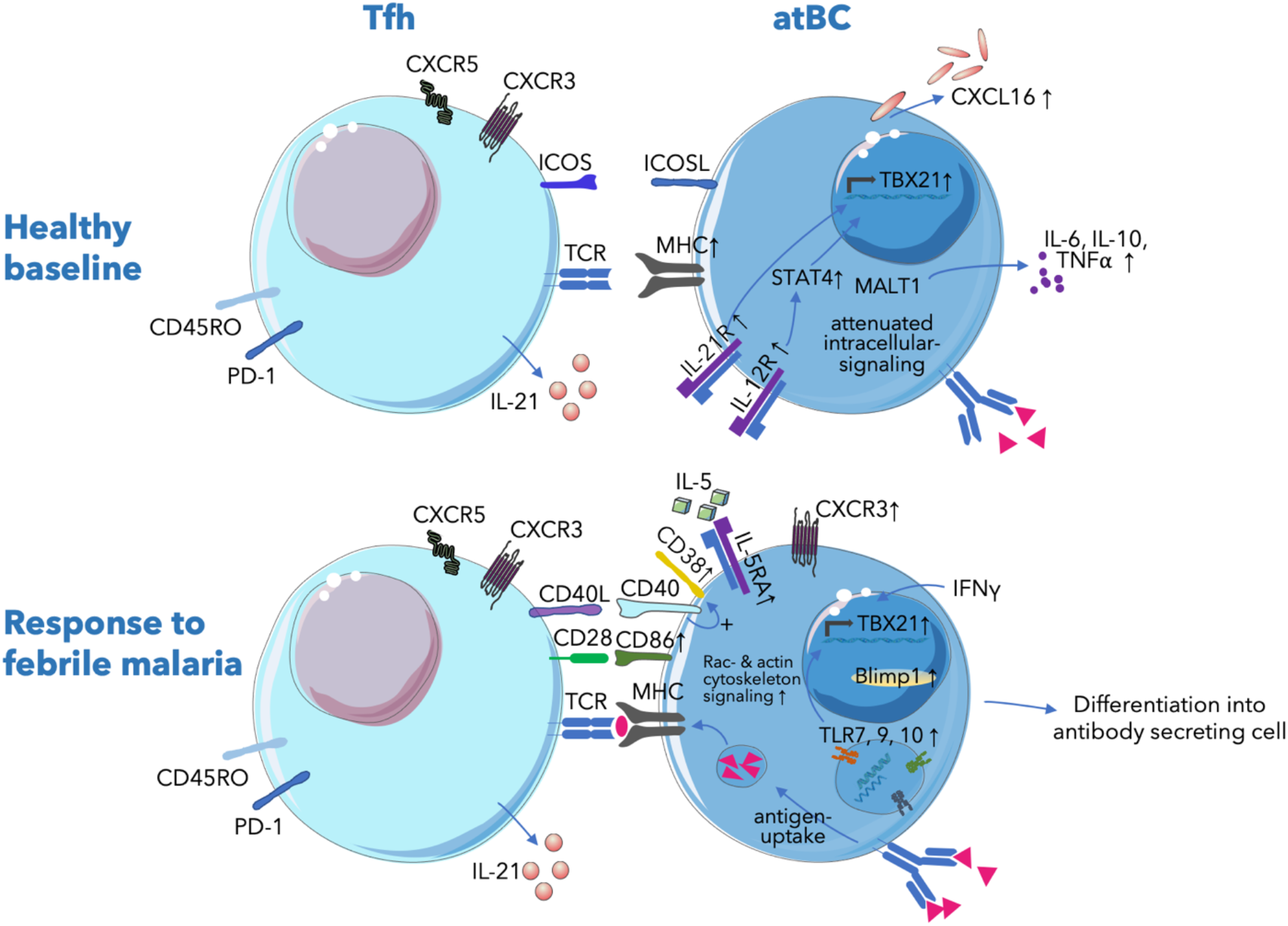
Model summarizing the observed phenotype of atBCs at baseline and after acute malaria and their potential interactions with cTfh cells. (**A**) At the healthy baseline, atBCs express ICOSL, MHC and the IL-21 receptor, suggesting that they are poised to interact with and receive signals from Tfh cells during subsequent infection. IL-21 signaling may contribute both directly and through STAT4 to the upregulation of T-bet. Further indicative of B-T cell interactions, *CXCL16*, a transmembrane chemokine typically expressed by antigen-presenting DCs in the T cell zone, is upregulated in atBCs. MALT1 is predicted to be linked to the upregulation of IL-10, IL-6 and TNF. (**B**) In response to acute malaria, atBCs upregulate CXCR3, which may facilitate migration to secondary lymphoid tissues. Upregulation of the co-stimulatory molecule CD86 and increased Rac- and actin cytoskeleton signaling, as well as increased signaling downstream of CD40 indicates interaction with GC Tfh cells. In addition, atBCs may capture membrane-bound antigen for presentation to Tfh cells (*34*). Acute malaria may also prime atBCs to respond to TLR7, 9 and 10 agonists, which together with IFNγ may contribute to T-bet expression. Upregulation of PRDM1 (Blimp1), a regulator of plasma cell differentiation, and CD38, an activation and plasma cell marker, suggest that atBCs may differentiate into antibody secreting cells during malaria.

At the healthy baseline *Pf*-specific atBCs also expressed lower levels of *TLR7, TLR9* and *TLR10* relative to actBCs and MBCs. Downregulation of *TLR9* by *Pf*-specific atBCs (i.e. *Pf*AMA1/MSP1-specific atBCs) in Malian donors is consistent with the finding that the TLR9-agonist CpG as an adjuvant did not enhance antibody or MBC responses to the *Pf*AMA1 or *Pf*MSP1 malaria vaccine candidates in Malian adults (*41*), while CpG enhanced responses to the same vaccines in malaria-naïve U.S. adults (*42*).

In response to acute febrile malaria, we found that both *Pf-*specific atBCs and actBCs were activated, proliferated, upregulated molecules involved in T cell interactions, and generally shared similar transcriptional profiles. For example, by flow cytometry we found that *Pf-*specific atBCs and actBCs upregulated Ki67, while RNA-seq analysis suggests that both subsets upregulate *CD38* and activate several intracellular pathways, including Rac-, CD40-, and actin cytoskeleton-signaling, as well as inositol phosphate compound pathways. Although *Pf-*specific atBCs and actBCs shared the putative upstream regulators IL-4, IL-5 and NUPR1, several other upstream regulators distinguished the two subsets, suggesting that different signals might activate the subsets during acute malaria. In response to malaria, the atBC markers T-bet and FcRL5 were upregulated in *Pf*-specific atBCs and actBCs, in line with a previous study of total human B cell subsets after malaria exposure (*24*) and studies showing that T-bet^hi^ HA-specific B cells arise following vaccination (*25, 43*). This indicates that T-bet and FcRL5 are stable markers of atBCs, as well as markers of recent activation of non-atBCs. However, by the end of the subsequent 6-month dry season (a period of negligible malaria transmission) T-bet and FcRL5 expression only decreased significantly in actBCs, suggesting heterogeneity in commitment to the T-bet^+^ FcRL5^+^ atBC phenotype in the absence of ongoing *Pf* exposure.

Relative to the healthy baseline, only *Pf*-specific atBCs upregulated *TLR7, TLR9* and *TLR10* expression in response to acute malaria. A previous study reported that the *Pf* erythrocyte membrane protein 1 (*Pf*EMP1) induces *TLR7* and *TLR10* expression and sensitizes B cells to TLR9 signaling (*44*). TLR9 signaling may in turn contribute to T-bet expression in *Pf-*specific atBCs (see model in Fig. 6), as it was reported that CpG or *Plasmodium* DNA together with IFN-γ induce T-bet expression in murine B cells (*45, 46*) and in our Ingenuity upstream regulator analysis, CpG was identified as a predicted upstream regulator in *Pf*-specific atBCs in response to malaria (Fig. S3).

In prior work we showed that malaria-associated atBCs have a markedly reduced capacity to be stimulated by soluble antigen through their BCR to proliferate, secrete cytokines or produce antibodies (*12*). A recent study found that malaria-associated atBCs signal through the BCR in response to planar lipid bilayers (PLB) containing F(ab′)_2_anti-λ/κ as a surrogate antigen to mimic presentation of antigen to B cells by follicular dendritic cells (*34*). The study also showed that atBCs capture and internalize anti-λ/κ bound to flexible plasma membrane sheets; and moreover, in response to PLB-associated anti-λ/κ, CpG, BAFF, IL-21, IL-2, and IL-10, atBCs expressed IRF4 mRNA, an early marker of plasma cell differentiation (*34*). However, it remained unknown if malaria-associated atBCs have the capacity to differentiate into ASCs.

*Ex vivo* RNA-seq analysis following acute malaria revealed that *Pf*-specific atBCs upregulate *PRDM1* (BLIMP-1) expression, an important regulator of plasma cell differentiation, as well as CD38, an activation and plasma cell marker. In response to acute malaria *Pf*-specific atBCs also upregulate the expression of molecules that mediate B-T cell interactions, including ICOSL, CD86, CD40, and MHC class II molecules. Together, these data led us to hypothesize that Tfh cells activate atBCs and drive them to differentiate into ASCs. Indeed, we found that in the presence of autologous Tfh cells and SEB, atBCs isolated from malaria-exposed individuals upregulate CD38 expression and differentiate into ASCs. Previous studies showed that frequencies of atBCs and circulating Tfh cells correlate with each other, and that malaria-induced IFN, as well as IL-21 and CD40L signals contribute to atBC differentiation (*8, 47*). Our data suggest that Tfh cells may also contribute to the activation and differentiation of atBCs into ASCs.

In addition to a potential role for atBCs in antibody production, we found evidence that atBCs may have the capacity to present antigen *in vivo*, consistent with the previous *in vitro* observation of internalization of membrane-associated antigen by atBCs (*34*). For example, we found by RNA-seq and flow cytometry that *Pf*-specific atBCs upregulate the expression of MHC class II molecules relative to MBCs. Additionally, atBCs express the integrin CD11c, which is also expressed by dendritic cells (DCs), and in mice, a CD11c^+^ subset phenotypically similar to atBCs was shown to have efficient antigen-presentation capacity by forming stable interactions with T cells (*48*). Furthermore, we found by RNA-seq that atBCs upregulate *CXCL16*, a transmembrane chemokine typically expressed by antigen-presenting DCs in the T cell zone.

This study has limitations. First, we investigated B cells specific for the immunodominant and polymorphic asexual blood-stage antigens *Pf*AMA1 and *Pf*MSP1. It will of interest to learn if the frequency and quality of *Pf*AMA1- and *Pf*MSP1-specific B cells are representative of B cells specific for other *Pf* antigens expressed during the various stages of the parasite life cycle. Second, we found that bulk atBCs can be activated to secrete antibodies by Tfh cells *in vitro*, and although *ex vivo* flow cytometry and RNA-seq data suggest that *Pf*-specific atBCs could also be activated by Tfh cells to secrete antibodies, the low frequency of *Pf*-specific atBCs precluded a direct test of this hypothesis *in vitro*. Third, *in vitro* activation by SEB bypasses the BCR and it is possible that a natural activation starting with BCR-stimulation and antigen-internalization would lead to an attenuation of responses to the immunological synapse between Tfh and B cell. Fourth, we previously found in the Mali cohort that acute malaria induces a Th1 cytokine response and activates a less-functional Th1-like CXCR3^+^ Tfh subset (*37*). In the current study, due to limited sample availability we were unable to test Tfh subsets of interest for their ability to activate atBCs and drive their differentiation to ASCs.

In summary, we found that in response to acute malaria *Pf*-specific atBCs are activated, increase in frequency, and upregulate molecules that mediate B-T cell interactions. Consistent with these *ex vivo* findings, we found that atBCs differentiate into ASCs when co-cultured with autologous Tfh cells from malaria-exposed individuals, together suggesting that atBCs may actively contribute to humoral immunity to infectious pathogens.

## MATERIALS AND METHODS

### Study design and ethical approval

The objective of this study was to examine pathogen-specific atBCs in the context of malaria infection, focusing on molecular changes indicative of atBC function by analyzing their phenotype, transcriptome and BCR sequence *ex vivo*. Samples and clinical data were obtained between 2015 and 2019 in a cohort study in Mali that has been described previously (*15*) and was conducted in Kalifabougou, Mali, where intense *P. falciparum* transmission occurs from June through December. Blood samples (8 mL) were collected by venipuncture from donors at the beginning (December) and end (May; healthy baseline) of each dry season, as well as 7 days after treatment of the first acute malaria episode (convalescence) of the ensuing 6-month malaria season. Acute malaria was defined by an axillary temperature of ≥37.5 °C or self-reported fever within 24 h and ≥2500 asexual parasites/μL of blood and no other cause of fever discernible by physical exam. Malaria episodes were treated according to Malian national guidelines with a 3-d standard course of artemether and lumefantrine.

The Ethics Committee of the Faculty of Medicine, Pharmacy, and Dentistry at the University of Sciences, Technique, and Technology of Bamako, and the Institutional Review Board of the National Institute of Allergy and Infectious Diseases, National Institutes of Health, approved this study. Written informed consent was obtained from adult participants and from the parents/guardians of participating children. The study is registered at ClinicalTrials.gov (NCT01322581). Blood samples from deidentified healthy U.S. adult donors were collected for research purposes at the National Institutes of Health Blood Bank under a National Institutes of Health institutional review board– approved protocol with informed consent (ClinicalTrials.gov NCT00001846).

### Flow cytometry with *Pf*MSP1, *Pf*AMA1- and HA-specific B cell probes

The full-length ectodomain of *Pf*AMA1 (strain FVO) and the C-terminal 42-kD region of *Pf*MSP1 (strain FUP) were expressed in Pichia pastoris and Escherichia coli, respectively, as previously described (*49, 50*). HA probes of two serotypes of influenza virus type A (H1N1 [A/California/04/2009] and H3N2 [A/Texas/50/ 2012]), consisting of the extracellular domain of HA, C-terminally fused to the trimeric FoldOn of T4 fibritin and the biotinylatable AviTag sequence, were expressed as previously described (*51*). Biotinylation, tetramerization, PBMC thawing and usage of B cell probes in flow cytometry have been described in detail elsewhere (*28*). Using a 1:1 ratio of biotin to protein, *Pf*AMA1 and *Pf*MSP1 were biotinylated using EZ-link Sulfo-NHS-LC-Biotinylation kit (Thermo Fisher Scientific) and HA proteins were biotinylated using AviTag technology (Avidity) according to published protocols (*51, 52*). Proteins were tetramerized with streptavidin-APC (Prozyme), streptavidin–Alexa Fluor 488 (Invitrogen), streptavidin-BV650 or streptavidin-BV785 (BD Biosciences). For flow cytometric analysis, cryopreserved PBMCs were thawed and stained with 100 ng of each B cell probe per sample, together with panels using the following labeled mAbs: CD3-BV510 (clone UCHT1), CD4-BV510 (clone SK3), CD8-BV510 (RPA-T8), CD14-BV510 (clone M5E2), CD16-BV510 (clone 3G8), CD56-BV510 (clone HCD56), CD10-APC-Cy7 or CD10-BV510 (clone Hi10a), CD19-ECD (clone J3-119), CD21-PE-Cy7 or CD21-BUV395 (clone B-ly4), CD27-BV605 (clone M-T271), IgD-BUV737 (clone IA-2), IgM-BUV395 (clone G20-127), IgG-Alexa Fluor 700 (clone G18-145), CD71-APC-Cy7 (clone CY1G4), CD86-PE (clone IT2.2), HLA-DR-APC-Cy7 (clone L243), Ki67-PE-Cy7 or KI67–Alexa Fluor 700 (clone 20Raj1), ICOSL-PE (clone 9F.8A4), and T-bet–PE (clone 4B10), FcRL5-PE (clone 509f6). Aqua dead cell stain was added for live/dead discrimination (Thermo Fisher Scientific). Stained samples were run on a BD Fortessa X20 (BD Biosciences), and data were analyzed using FlowJo (FlowJo Software).

### Single-cell BCR amplification and sequence analysis

For BCR sequencing, individual CD19^+^ CD10^−^ *Pf*^+^ were single-cell sorted into 96-well plates using a FACSAria II (BD Biosciences). Index sorting was used to determine B cell subsets by CD21 and CD27 expression. BCR sequencing was performed as previously described (*25, 28, 53*). Briefly, cDNA was synthesized from sorted cells using random hexamers and Superscript III RT (Thermo Fisher Scientific). Ig heavy chain genes were then PCR amplified separately with two rounds of nested PCR using DreamTaq Mastermix (Thermo Fisher Scientific). PCR products were Sanger sequenced by GenScript, and sequences were analyzed using IMGT/V-QUEST (*32*).

### Transcriptomic analysis using SMART-Seq methodology

#### Sample preparation, cDNA synthesis and library Preparation

Yielding 12 samples per donor, *Pf*AMA1/PfMSP1- and HA-specific atBC, actBC, and MBC populations from May and convalescence time points (donor ages 11-14 years) were sorted using a FACSAria IIIu (BD Biosciences) directly into 1.7 ml eppendorf tubes containing 12.0 μl 1X reaction buffer (Takara) for cDNA synthesis using the SMART-Seq v4 Ultra low Input RNA kit (Takara) following the manufacturer’s protocol. Sample processing was performed in 96-well plates and to reduce plate position effects, samples were arranged in a balanced and randomized order. Flow-sorted cell counts ranged from 1-47 (see Table S1) and to normalize, a volume of lysate corresponding to 4 cells was used as input for library construction from each sample. cDNA was assessed and quantified using the Bioanalyzer 2100 (Agilent) and per sample 5 ng was used as template to generate sequencing libraries. cDNA was brought to a final volume of 15 μl and sheared using a LE220 focused-ultrasonicator (Covaris, Woburn, MA) with the following shearing parameters: peak power 180; duty factor 10; cycles/burst 50; 19°C; 270 seconds. The SMARTer ThruPLEX DNA-Seq kit (Takara, Rockville, MD) was utilized to generate libraries from 10 μl of sheared cDNA template, following the manufacturer’s instructions with a total of 11 PCR cycles. Purified libraries were assessed on the Bioanalyzer 2100 (Agilent Technologies, Santa Clara, CA), leading to the exclusion of two samples with low library yield.

#### Illumina sequencing

Two microliters of each purified ThruPLEX library were combined to create a library pool which was then quantified on the CFX96 Touch real-time PCR instrument (BioRad, Hercules, CA) using the Kapa Library Quant Universal qPCR mix and kit instructions (Kapa Biosystems, Wilmington, MA). The library pool was normalized to a 2 nM concentration, titrated to 10.5 pM, and sequenced as 2 × 150 bp reads using the MiSeq V2 Nano kit, according to the manufacturer’s instructions (Illumina, San Diego, CA). The read counts from the MiSeq nano run were used to create a new pool of each library containing equimolar amounts of each sample for the final instrument runs. The library pool containing adjusted library amounts was quantified using the Kapa Library Quant Universal kit and normalized to 2 nM. The pool was titrated to 1.8 -1.9 pM and sequenced as 2 × 75 bp reads in a total of nine runs on the NextSeq 550 instrument using the high output v2.5 kit (Illumina, San Diego, CA).

#### Bioinformatic processing

Following sequencing, raw files were converted to fastq files using bcl2fastq (v2.20.0.422, Illumina Inc. San Diego, CA), trimmed of adapter sequences using cutadapt version 1.12 (*54*) and quality-trimmed and filtered using fastq_quality_trimmer and fastq_quality_filter tools from the FASTX Toolkit, v 0.0.14 (*55*). Singletons were removed and trimmed reads were sorted in coordinate order using a custom Perl script. Quality trimmed and sorted reads were aligned to the human genome reference build GRCh38 using HISAT, version 2.1.0 (*56*). Sequence Alignment/MAP (SAM) files were converted to Binary Alignment/Map (BAM) format, sorted and indexed using SAMTOOLS version 1.8 (*57, 58*). The number of reads per gene locus was obtained from the aligned BAM files using HTSeq-count, version 0.9.1 (*58*) and a counts matrix was generated using a custom Perl script.

#### Quantification and statistical analysis

NGS transcriptome libraries were generated from 73 healthy baseline samples and 95 convalescence samples using the SMART-Seq v4 Ultra low Input RNA kit (Takara), an optimized version of the SMART-Seq2 method. Eight healthy baseline libraries and four convalescence libraries were excluded due to poor quality and 156 libraries were sequenced on the Illumina platform. Using the counts matrix, read data was normalized and analyzed for differential expression with the Bioconductor package DESeq2, v 1.30.0 (*59*). Additional low-expression filters were applied to remove genes with zero counts in greater than 140 samples or <158 total read counts across all 156 samples and to remove samples with zero counts for >90% of genes, resulting in the removal of 11,433 genes. Principal component analysis was performed in R using the “plotPCA()” function from the “affycoretools” and samples identified as “outlier” were removed. The above filters removed 20 healthy baseline samples and 16 convalescence samples. 45 samples at healthy baseline and 75 samples at convalescence remained and were used for DEG analysis. Differential expression analysis and statistical analysis of comparisons were performed using custom contrasts in DESeq2 (BioConductor version 1.30.0) with R software (version 4.0.3). Top 250 DEGs for each subsets were identified and principal component analysis plot were generated in R using the “plotPCA()” function from the “affycoretools” package library. Heatmaps were generated using GraphPad Prism. Apart from heatmaps, where all significant DEGs were indicated by *, abbreviations for p values in all other graphs were as follows: p<0.05 = ^*^, p<0.01 = ^**^, p<0.001 = ^***^, p<0.0001 = ^****^.

#### Ingenuity Pathway Analysis (IPA)

IPA was performed using DEGs with a cut-off FDR of <0.2, with no fold change (FC) cut-off. Predicted upstream regulators were selected based on absolute z-score of >2 and p value <0.005. Significant overlap beween DEGs and canonical pathways were determined by the Fisher’s exact test using a cut-off Benjamini-Hochberg adjusted p<0.05.

### Co-culture of Tfh cells and B cell subsets

PBMCs were FACS sorted into PD-1^+^ CXCR5^+^ CD45RO^+^ CD3^+^ CD4^+^ Tfh cells, CD19^+^ CD10-CD21^-^ CD27^-^ atBCs, CD19^+^ CD10-CD21^+^ CD27^+^ MBCs and CD19^+^ CD10^−^CD21^-^ CD27^+^ actBCs. Antibody clones: CD3-PE-Cy5 (clone UCHT1), CD4-BV421 (clone SK3), CD10-PE (clone Hi10a), CD19-ECD (clone J3-119), CD21-PE-Cy7 (clone B-ly4), CD27-BV605 (clone M-T271), CD45RO-BV785 (clone UCHL1), CXCR5-AF488 (clone RF8B2), PD-1-APC (clone EH12.2H7). 18.000 – 25.000 cells of each B cell subset were co-cultured with Tfh cells at a 1:1 ratio for 7 days in complete medium with SEB (1.5 μg/ml; Sigma-Aldrich). After co-culture, CD38 surface expression was assessed by flow cytometry (CD38-APC, clone HIT1) and Ig levels in supernatants were measured with ProcartaPlex Human Antibody 7plex Isotyping Panel (eBioscience).

## Acknowledgments

We thank the residents of Kalifabougou, Mali, for participating in this study. *Pf*AMA1 and *Pf*MSP1 proteins were kindly provided by David Narum at the Laboratory of Malaria Immunology and Vaccinology, National Institute of Allergy and Infectious Diseases, NIH. HA proteins were kindly shared by Sarah Andrews, Michael J. Chambers and Adrian McDermott at the Vaccine Research Center, National Institute of Allergy and Infectious Diseases, NIH. We also thank the NIAID and LIG flow cytometry facilities (Ludmila Krymskaya and Calvin Eigsti). This work was supported by the Division of Intramural Research, National Institute of Allergy and Infectious Diseases, National Institutes of Health.

## Author contributions

Conceptualization: CSH, PDC

Methodology: CSH, SLA, MEP, SL

Investigation: CSH, JS, SLA, CT

Consulting Statistician: JS, SLA

Visualization: CSH

Funding acquisition: PDC

Project administration: SD, KK, AO, CM, BT, PDC

Supervision: KK, BT, PDC, CSH

Writing – original draft: CSH

Writing – review & editing: CSH, SLA, JS, PDC

## Data availability

The BCR sequencing data were deposited into the National Center for Biotechnology Information and are accessible via BioProject accession no. PRJNA769078 (reviewer link: https://dataview.ncbi.nlm.nih.gov/object/PRJNA769078?reviewer=r2dd7qnlf3makouqf0t6qc21iv). The RNAseq data discussed in this publication have been deposited in NCBI’s Gene Expression Omnibus (GEO)(*60*) and are accessible through GEO Series accession number GSE183744 (https://www.ncbi.nlm.nih.gov/geo/query/acc.cgi?acc=GSE183744).

## REFERENCES

1. A. E. Braddom, G. Batugedara, S. Bol, E. M. Bunnik, Potential functions of atypical memory B cells in Plasmodium-exposed individuals, Int. J. Parasitol. 50, 1033–1042 (2020).

2. J. L. Karnell, V. Kumar, J. Wang, S. Wang, E. Voynova, R. Ettinger, Role of CD11c+ T-bet+ B cells in human health and disease, Cell. Immunol. 321, 40–45 (2017).

3. N. H. Wildner, P. Ahmadi, S. Schulte, F. Brauneck, M. Kohsar, M. Lütgehetmann, C. Beisel, M. M. Addo, F. Haag, J. Schulze Zur Wiesch, B cell analysis in SARS-CoV-2 versus malaria: Increased frequencies of plasmablasts and atypical memory B cells in COVID-19, J. Leukoc. Biol. 109, 77–90 (2021).

4. S. A. Joosten, K. E. van Meijgaarden, F. Del Nonno, A. Baiocchini, L. Petrone, V. Vanini, H. H. Smits, F. Palmieri, D. Goletti, T. H. M. Ottenhoff, D. M. Lewinsohn, Ed. Patients with Tuberculosis Have a Dysfunctional Circulating B-Cell Compartment, Which Normalizes following Successful Treatment, PLoS Pathog. 12, e1005687 (2016).

5. S. Moir, J. Ho, A. Malaspina, W. Wang, A. C. DiPoto, M. A. O’Shea, G. Roby, S. Kottilil, J. Arthos, M.A. Proschan, T.-W. Chun, A. S. Fauci, Evidence for HIV-associated B cell exhaustion in a dysfunctional memory B cell compartment in HIV-infected viremic individuals, J. Exp. Med. 205, 1797–1805 (2008).

6. G. E. Weiss, P. D. Crompton, S. Li, L. A. Walsh, S. Moir, B. Traore, K. Kayentao, A. Ongoiba, O. K. Doumbo, S. K. Pierce, Atypical memory B cells are greatly expanded in individuals living in a malaria-endemic area, The Journal of Immunology 183, 2176–2182 (2009).

7. S. Portugal, N. Obeng-Adjei, S. Moir, P. D. Crompton, S. K. Pierce, Atypical memory B cells in human chronic infectious diseases: An interim report, Cell. Immunol. 321, 18–25 (2017).

8. N. Obeng-Adjei, S. Portugal, P. Holla, S. Li, H. Sohn, A. Ambegaonkar, J. Skinner, G. Bowyer, O. K. Doumbo, B. Traore, S. K. Pierce, P. D. Crompton,Malaria-induced interferon-γ drives the expansion of Tbethi atypical memory B cells, PLoS Pathog. 13, e1006576 (2017).

9. P. Holla, B. Dizon, A. A. Ambegaonkar, N. Rogel, E. Goldschmidt, A. K. Boddapati, H. Sohn, D. Sturdevant, J. W. Austin, L. Kardava, L. Yuesheng, P. Liu, S. Moir, S. K. Pierce, A. Madi, Shared transcriptional profiles of atypical B cells suggest common drivers of expansion and function in malaria, HIV, and autoimmunity, Sci Adv 7, eabg8384 (2021).

10. H. J. Sutton, R. Aye, A. H. Idris, R. Vistein, E. Nduati, O. Kai, J. Mwacharo, X. Li, X. Gao, T. D. Andrews, M. Koutsakos, T. H. O. Nguyen, M. Nekrasov, P. Milburn, A. Eltahla, A. A. Berry, N. Kc, S. Chakravarty, B. K. L. Sim, A. K. Wheatley, S. J. Kent, S. L. Hoffman, K. E. Lyke, P. Bejon, F. Luciani, K. Kedzierska, R. A. Seder, F. M. Ndungu, I. A. Cockburn, Atypical B cells are part of an alternative lineage of B cells that participates in responses to vaccination and infection in humans, Cell Rep 34, 108684 (2021).

11. R. T. Sullivan, C. C. Kim, M. F. Fontana, M. E. Feeney, P. Jagannathan, M. J. Boyle, C. J. Drakeley, I. Ssewanyana, F. Nankya, H. Mayanja-Kizza, G. Dorsey, B. Greenhouse, FCRL5 Delineates Functionally Impaired Memory B Cells Associated with Plasmodium falciparum Exposure, PLoS Pathog. 11, e1004894 (2015).

12. S. Portugal, C. M. Tipton, H. Sohn, Y. Kone, J. Wang, S. Li, J. Skinner, K. Virtaneva, D. E. Sturdevant, S. F. Porcella, O. K. Doumbo, S. Doumbo, K. Kayentao, A. Ongoiba, B. Traore, I. Sanz, S. K. Pierce, P. D. Crompton, Malaria-associated atypical memory B cells exhibit markedly reduced B cell receptor signaling and effector function, Elife 4 (2015), doi:10.7554/eLife.07218.

13. World Health Organization, World malaria report 2020 -20 years of global progress & challenges,, 1–300 (2020).

14. J. Langhorne, F. M. Ndungu, A.-M. Sponaas, K. Marsh, Immunity to malaria: more questions than answers, Nature Publishing Group 9, 725–732 (2008).

15. T. M. Tran, S. Li, S. Doumbo, D. Doumtabe, C.-Y. Huang, S. Dia, A. Bathily, J. Sangala, Y. Kone, A. Traore, M. Niangaly, C. Dara, K. Kayentao, A. Ongoiba, O. K. Doumbo, B. Traore, P. D. Crompton, An intensive longitudinal cohort study of Malian children and adults reveals no evidence of acquired immunity to Plasmodium falciparum infection, Clin. Infect. Dis. 57, 40–47 (2013).

16. J. Tan, H. Cho, T. Pholcharee, L. S. Pereira, S. Doumbo, D. Doumtabe, B. J. Flynn, A. Schön, S. Kanatani, S. O. Aylor, D. Oyen, R. Vistein, L. Wang, M. Dillon, J. Skinner, M. Peterson, S. Li, A. H. Idris, A. Molina-Cruz, M. Zhao, L. R. Olano, P. J. Lee, A. Roth, P. Sinnis, C. Barillas-Mury, K. Kayentao, A. Ongoiba, J. R. Francica, B. Traore, I. A. Wilson, R. A. Seder, P. D. Crompton, Functional human IgA targets a conserved site on malaria sporozoites, Sci Transl Med 13 (2021), doi:10.1126/scitranslmed.abg2344.

17. S. Cohen, I. A. McGregor, S. Carrington, Gamma-globulin and acquired immunity to human malaria, Nature 192, 733–737 (1961).

18. G. E. Weiss, E. H. Clark, S. Li, B. Traore, K. Kayentao, A. Ongoiba, J. N. Hernandez, O. K. Doumbo, S. K. Pierce, O. H. Branch, P. D. Crompton, A positive correlation between atypical memory B cells and Plasmodium falciparum transmission intensity in cross-sectional studies in Peru and Mali, PLoS ONE 6, e15983 (2011).

19. S. I. Nogaro, J. C. Hafalla, B. Walther, E. J. Remarque, K. K. A. Tetteh, D. J. Conway, E. M. Riley, M. Walther, The breadth, but not the magnitude, of circulating memory B cell responses to P. falciparum increases with age/exposure in an area of low transmission, PLoS ONE 6, e25582 (2011).

20. J. Illingworth, N. S. Butler, S. Roetynck, J. Mwacharo, S. K. Pierce, P. Bejon, P. D. Crompton, K. Marsh, F. M. Ndungu, Chronic exposure to Plasmodium falciparum is associated with phenotypic evidence of B and T cell exhaustion, J. Immunol. 190, 1038–1047 (2013).

21. M. F. Muellenbeck, B. Ueberheide, B. Amulic, A. Epp, D. Fenyo, C. E. Busse, M. Esen, M. Theisen, B. Mordmüller, H. Wardemann, Atypical and classical memory B cells produce Plasmodium falciparum neutralizing antibodies, J. Exp. Med. 210, 389–399 (2013).

22. R. Aye, H. J. Sutton, E. W. Nduati, O. Kai, J. Mwacharo, J. Musyoki, E. Otieno, J. Wambua, P. Bejon, I. A. Cockburn, F. M. Ndungu, Malaria exposure drives both cognate and bystander human B cells to adopt an atypical phenotype, Eur. J. Immunol. 50, 1187–1194 (2020).

23. D. Perez-Mazliah, P. J. Gardner, E. Schweighoffer, S. McLaughlin, C. Hosking, I. Tumwine, R. S. Davis, A. J. Potocnik, V. L. Tybulewicz, J. Langhorne, Plasmodium-specific atypical memory B cells are short-lived activated B cells, Elife 7, 435 (2018).

24. C. Sundling, C. Rönnberg, V. Yman, M. Asghar, P. Jahnmatz, T. Lakshmikanth, Y. Chen, J. Mikes, M. N. Forsell, K. Sondén, A. Achour, P. Brodin, K. E. Persson, A. Färnert, B cell profiling in malaria reveals expansion and remodelling of CD11c+ B cell subsets, JCI Insight 5, 633 (2019).

25. S. F. Andrews, M. J. Chambers, C. A. Schramm, J. Plyler, J. E. Raab, M. Kanekiyo, R. A. Gillespie, A. Ransier, S. Darko, J. Hu, X. Chen, H. M. Yassine, J. C. Boyington, M. C. Crank, G. L. Chen, E. Coates, J. R. Mascola, D. C. Douek, B. S. Graham, J. E. Ledgerwood, A. B. McDermott, Activation Dynamics and Immunoglobulin Evolution of Pre-existing and Newly Generated Human Memory B cell Responses to Influenza Hemagglutinin, Immunity (2019), doi:10.1016/j.immuni.2019.06.024.

26. C. C. Kim, A. M. Baccarella, A. Bayat, M. Pepper, M. F. Fontana, FCRL5+ Memory B Cells Exhibit Robust Recall Responses, Cell Rep 27, 1446–1460.e4 (2019).

27. A. H. Ellebedy, K. J. L. Jackson, H. T. Kissick, H. I. Nakaya, C. W. Davis, K. M. Roskin, A. K. McElroy, C. M. Oshansky, R. Elbein, S. Thomas, G. M. Lyon, C. F. Spiropoulou, A. K. Mehta, P. G. Thomas, S. D. Boyd, R. Ahmed, Defining antigen-specific plasmablast and memory B cell subsets in human blood after viral infection or vaccination, Nature Publishing Group 17, 1226–1234 (2016).

28. C. S. Hopp, P. Sekar, A. Diouf, K. Miura, K. Boswell, J. Skinner, C. M. Tipton, M. E. Peterson, M. J. Chambers, S. Andrews, J. Lu, J. Tan, S. Li, S. Doumbo, K. Kayentao, A. Ongoiba, B. Traore, S. Portugal, P. D. Sun, C. Long, R. A. Koup, E. O. Long, A. B. McDermott, P. D. Crompton, Plasmodium falciparum-specific IgM B cells dominate in children, expand with malaria, and produce functional IgM, J. Exp. Med. 218 (2021), doi:10.1084/jem.20200901.

29. N. Talla Nzussouo, J. Duque, A. A. Adedeji, D. Coulibaly, S. Sow, Z. Tarnagda, I. Maman, A. Lagare, S. Makaya, M. B. Elkory, H. Kadjo Adje, P. A. Shilo, B. Tamboura, A. Cisse, K. Badziklou, H. B. Maïnassara, A. O. Bara, A. M. Keita, T. Williams, A. Moen, M.-A. Widdowson, M. McMorrow, Epidemiology of influenza in West Africa after the 2009 influenza A(H1N1) pandemic, 2010-2012, BMC Infect. Dis. 17, 745–8 (2017).

30. C. M. Andrade, H. Fleckenstein, R. Thomson-Luque, S. Doumbo, N. F. Lima, C. Anderson, J. Hibbert, C. S. Hopp, T. M. Tran, S. Li, M. Niangaly, H. Cisse, D. Doumtabe, J. Skinner, D. Sturdevant, S. Ricklefs, K. Virtaneva, M. Asghar, M. V. Homann, L. Turner, J. Martins, E. L. Allman, M.-E. N’Dri, V. Winkler, M. Llinás, C. Lavazec, C. Martens, A. Färnert, K. Kayentao, A. Ongoiba, T. Lavstsen, N. S. Osório, T. D. Otto, M. Recker, B. Traore, P. D. Crompton, S. Portugal, Increased circulation time of Plasmodium falciparum underlies persistent asymptomatic infection in the dry season, Nat. Med. 157, 1137–12 (2020).

31. L. Kardava, S. Moir, N. Shah, W. Wang, R. Wilson, C. M. Buckner, B. H. Santich, L. J. Y. Kim, E. E. Spurlin, A. K. Nelson, A. K. Wheatley, C. J. Harvey, A. B. McDermott, K. W. Wucherpfennig, T.-W. Chun, J. S. Tsang, Y. Li, A. S. Fauci, Abnormal B cell memory subsets dominate HIV-specific responses in infected individuals, J. Clin. Invest. 124, 3252–3262 (2014).

32. V. Giudicelli, X. Brochet, M.-P. Lefranc, IMGT/V-QUEST: IMGT standardized analysis of the immunoglobulin (IG) and T cell receptor (TR) nucleotide sequences, Cold Spring Harb Protoc 2011, 695–715 (2011).

33. M. Masilamani, D. Kassahn, S. Mikkat, M. O. Glocker, H. Illges, B cell activation leads to shedding of complement receptor type II (CR2/CD21), Eur. J. Immunol. 33, 2391–2397 (2003).

34. A. A. Ambegaonkar, K. Kwak, H. Sohn, J. Manzella-Lapeira, J. Brzostowski, S. K. Pierce, Expression of inhibitory receptors by B cells in chronic human infectious diseases restricts responses to membrane-associated antigens, Sci Adv 6, eaba6493 (2020).

35. W. Song, J. Craft, T follicular helper cell heterogeneity: Time, space, and function, Immunol. Rev. 288, 85–96 (2019).

36. R. Morita, N. Schmitt, S. E. Bentebibel, R. Ranganathan, L. Bourdery, G. Zurawski, E. Foucat, M. Dullaers, S. Oh, N. Sabzghabaei, E. M. Lavecchio, M. Punaro, V. Pascual, J. Banchereau, H. Ueno, Human Blood CXCR5+CD4+ T Cells Are Counterparts of T Follicular Cells and Contain Specific Subsets that Differentially Support Antibody Secretion, Immunity 34, 108–121 (2011).

37. N. Obeng-Adjei, S. Portugal, T. M. Tran, T. B. Yazew, J. Skinner, S. Li, A. Jain, P. L. Felgner, O. K. Doumbo, K. Kayentao, A. Ongoiba, B. Traore, P. D. Crompton, Circulating Th1-Cell-type Tfh Cells that Exhibit Impaired B Cell Help Are Preferentially Activated during Acute Malaria in Children, Cell Rep 13, 425–439 (2015).

38. A. E. Braddom, S. Bol, S. J. Gonzales, R. A. Reyes, K. Musinguzi, F. Nankya, I. Ssewanyana, B. Greenhouse, E. M. Bunnik, B Cell Receptor Repertoire Analysis in Malaria-Naive and Malaria-Experienced Individuals Reveals Unique Characteristics of Atypical Memory B Cells, mSphere, e0072621 (2021).

39. D. Durali, M.-G. de Goër de Herve, J. Giron-Michel, B. Azzarone, J.-F. Delfraissy, Y. Taoufik, In human B cells, IL-12 triggers a cascade of molecular events similar to Th1 commitment, Blood 102, 4084–4089 (2003).

40. S. Wang, J. Wang, V. Kumar, J. L. Karnell, B. Naiman, P. S. Gross, S. Rahman, K. Zerrouki, R. Hanna, C. Morehouse, N. Holoweckyj, H. Liu, Autoimmunity Molecular Medicine Team, Z. Manna, R. Goldbach-Mansky, S. Hasni, R. Siegel, M. Sanjuan, K. Streicher, M. P. Cancro, R. Kolbeck, R. Ettinger, IL-21 drives expansion and plasma cell differentiation of autoreactive CD11chiT-bet+ B cells in SLE, Nat Commun 9, 1758–14 (2018).

41. B. Traore, Y. Kone, S. Doumbo, D. Doumtabe, A. Traore, P. D. Crompton, M. Mircetic, C.-Y. Huang, K. Kayentao, A. Dicko, I. Sagara, R. D. Ellis, K. Miura, A. Guindo, L. H. Miller, O. K. Doumbo, S. K. Pierce, The TLR9 agonist CpG fails to enhance the acquisition of Plasmodium falciparum-specific memory B cells in semi-immune adults in Mali, Vaccine 27, 7299–7303 (2009).

42. P. D. Crompton, M. Mircetic, G. Weiss, A. Baughman, C.-Y. Huang, D. J. Topham, J. J. Treanor, I. Sanz, F. E.-H. Lee, A. P. Durbin, K. Miura, D. L. Narum, R. D. Ellis, E. Malkin, G. E. D. Mullen, L. H. Miller, L. B. Martin, S. K. Pierce, The TLR9 ligand CpG promotes the acquisition of Plasmodium falciparum-specific memory B cells in malaria-naive individuals, The Journal of Immunology 182, 3318–3326 (2009).

43. D. Lau, L. Y.-L. Lan, S. F. Andrews, C. Henry, K. T. Rojas, K. E. Neu, M. Huang, Y. Huang, B. DeKosky, A.-K. E. Palm, G. C. Ippolito, G. Georgiou, P. C. Wilson, Low CD21 expression defines a population of recent germinal center graduates primed for plasma cell differentiation, Sci Immunol 2, eaai8153 (2017).

44. O. Simone, M. T. Bejarano, S. K. Pierce, S. Antonaci, M. Wahlgren, M. Troye-Blomberg, D. Donati, TLRs innate immunereceptors and Plasmodium falciparum erythrocyte membrane protein 1 (PfEMP1) CIDR1α-driven human polyclonal B-cell activation, Acta Trop. 119, 144–150 (2011).

45. N. Liu, N. Ohnishi, L. Ni, S. Akira, K. B. Bacon, CpG directly induces T-bet expression and inhibits IgG1 and IgE switching in B cells, Nat. Immunol. 4, 687–693 (2003).

46. J. Rivera-Correa, J. J. Guthmiller, R. Vijay, C. Fernandez-Arias, M. A. Pardo-Ruge, S. Gonzalez, N. S. Butler, A. Rodríguez, Plasmodium DNA-mediated TLR9 activation of T-bet(+) B cells contributes to autoimmune anaemia during malaria, Nat Commun 8, 1282 (2017).

47. B. Keller, V. Strohmeier, I. Harder, S. Unger, K. J. Payne, G. Andrieux, M. Boerries, P. T. Felixberger, J. J. M. Landry, A. Nieters, A. Rensing-Ehl, U. Salzer, N. Frede, S. Usadel, R. Elling, C. Speckmann, I. Hainmann, E. Ralph, K. Gilmour, M. W. J. Wentink, M. van der Burg, H. S. Kuehn, S. D. Rosenzweig, U. Kölsch, H. von Bernuth, P. Kaiser-Labusch, F. Gothe, S. Hambleton, A. D. Vlagea, A. Garcia Garcia, L. Alsina, G. Markelj, T. Avcin, J. Vasconcelos, M. Guedes, J.-Y. Ding, C.-L. Ku, B. Shadur, D. T. Avery, N. Venhoff, J. Thiel, H. Becker, L. Erazo-Borrás, C. M. Trujillo-Vargas, J. L. Franco, C. Fieschi, S. Okada, P. E. Gray, G. Uzel, J.-L. Casanova, M. Fliegauf, B. Grimbacher, H. Eibel, S. Ehl, R. E. Voll, M. Rizzi, P. Stepensky, V. Benes, C. S. Ma, C. Bossen, S. G. Tangye, K. Warnatz, The expansion of human T-bethighCD21low B cells is T cell dependent, Sci Immunol 6, eabh0891 (2021).

48. A. V. Rubtsov, K. Rubtsova, J. W. Kappler, J. Jacobelli, R. S. Friedman, P. Marrack, CD11c-Expressing B Cells Are Located at the T Cell/B Cell Border in Spleen and Are Potent APCs, The Journal of Immunology 195, 71–79 (2015).

49. R. L. Shimp, L. B. Martin, Y. Zhang, B. S. Henderson, P. Duggan, N. J. MacDonald, J. Lebowitz, A. Saul, D. L. Narum, Production and characterization of clinical grade Escherichia coli derived Plasmodium falciparum 42 kDa merozoite surface protein 1 (MSP1(42)) in the absence of an affinity tag, Protein Expr. Purif. 50, 58–67 (2006).

50. R. D. Ellis, Y. Wu, L. B. Martin, D. Shaffer, K. Miura, J. Aebig, A. Orcutt, K. Rausch, D. Zhu, A. Mogensen, M. P. Fay, D. L. Narum, C. Long, L. Miller, A. P. Durbin, T. L. Richie, Ed. Phase 1 Study in Malaria Naïve Adults of BSAM2/Alhydrogel®+CPG 7909, a Blood Stage Vaccine against P. falciparum Malaria, PLoS ONE 7, e46094–11 (2012).

51. J. R. R. Whittle, A. K. Wheatley, L. Wu, D. Lingwood, M. Kanekiyo, S. S. Ma, S. R. Narpala, H. M. Yassine, G. M. Frank, J. W. Yewdell, J. E. Ledgerwood, C.-J. Wei, A. B. McDermott, B. S. Graham, R. A. Koup, G. J. Nabel, Flow cytometry reveals that H5N1 vaccination elicits cross-reactive stem-directed antibodies from multiple Ig heavy-chain lineages, J. Virol. 88, 4047–4057 (2014).

52. J. J. Taylor, R. J. Martinez, P. J. Titcombe, L. O. Barsness, S. R. Thomas, N. Zhang, S. D. Katzman, M. K. Jenkins, D. L. Mueller, Deletion and anergy of polyclonal B cells specific for ubiquitous membrane-bound self-antigen, J. Exp. Med. 209, 2065–2077 (2012).

53. T. Tiller, E. Meffre, S. Yurasov, M. Tsuiji, M. C. Nussenzweig, H. Wardemann, Efficient generation of monoclonal antibodies from single human B cells by single cell RT-PCR and expression vector cloning, J. Immunol. Methods 329, 112–124 (2008).

54. M. Marcel, Cutadapt removes adapter sequences from high-throughput sequencing reads, EMBnet 17, 10–12 (2011).

55. G. Hannon, The FASTX-Toolkit (2010).

56. D. Kim, B. Langmead, S. L. Salzberg, HISAT: a fast spliced aligner with low memory requirements, Nat Meth 12, 357–360 (2015).

57. H. Li, B. Handsaker, A. Wysoker, T. Fennell, J. Ruan, N. Homer, G. Marth, G. Abecasis, R. Durbin, 1000 Genome Project Data Processing Subgroup, The Sequence Alignment/Map format and SAMtools, Bioinformatics 25, 2078–2079 (2009).

58. S. Anders, P. T. Pyl, W. Huber, HTSeq--a Python framework to work with high-throughput sequencing data, Bioinformatics 31, 166–169 (2015).

59. M. I. Love, W. Huber, S. Anders, Moderated estimation of fold change and dispersion for RNA-seq data with DESeq2, Genome Biol. 15, 550–21 (2014).

60. R. Edgar, M. Domrachev, A. E. Lash, Gene Expression Omnibus: NCBI gene expression and hybridization array data repository, Nucleic Acids Res. 30, 207–210 (2002).

