## Supplemental Material for "Atypical B cells upregulate co-stimulatory molecules during malaria and secrete antibodies with T follicular helper cell support"

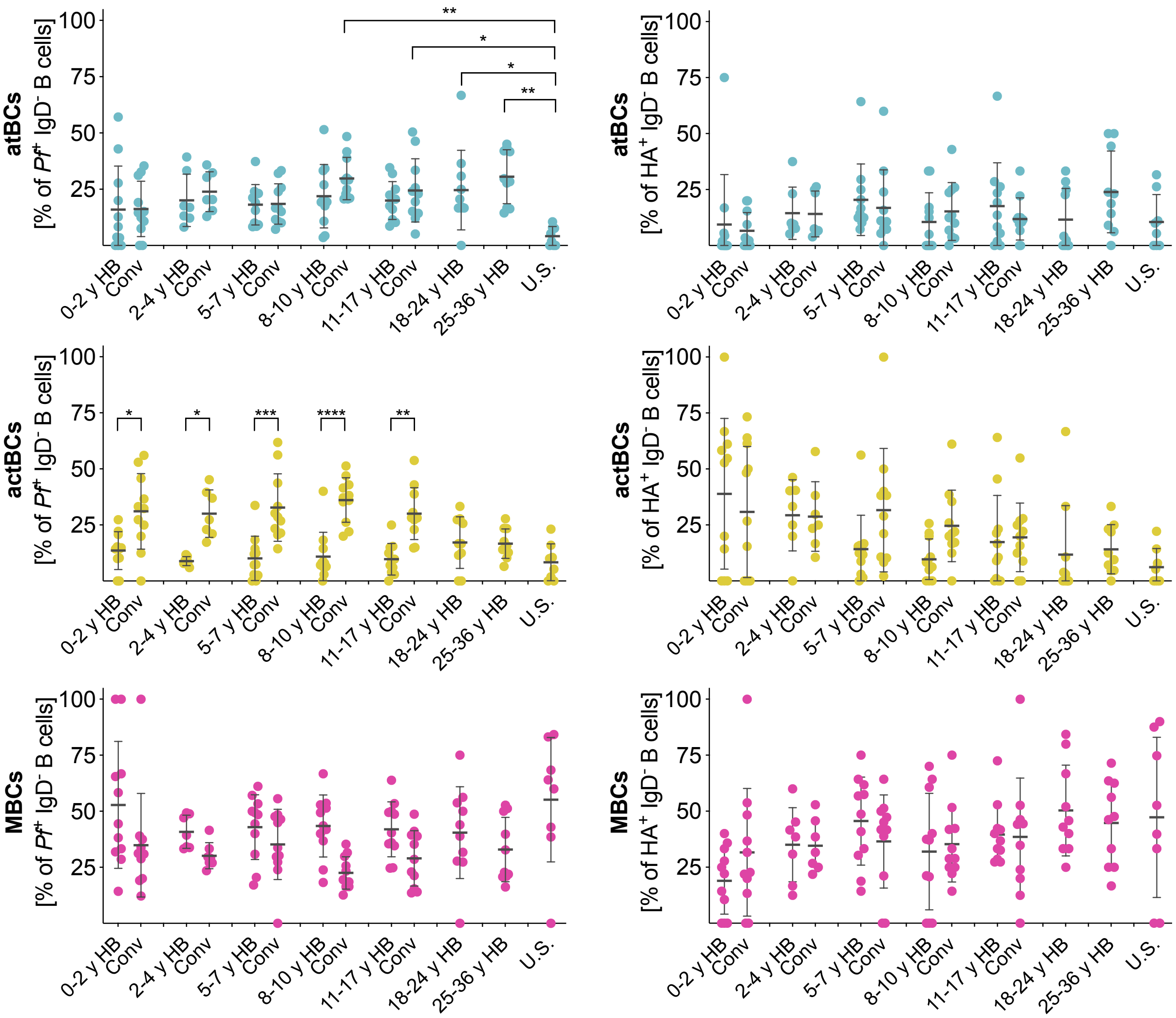


**Supplementary Figure 1. The B cell subset distribution of *Pf*- and HA-specific B cells is comparable in children and adults.** The percentage of B cell subsets (atBCs, top; actBCs, middle; MBCs, bottom) among *Pf*- and HA-specific IgD$^-$ B cells at the healthy baseline (HB) and one week after treatment of febrile malaria (Conv). Mean percentages and SDs are shown for Malian individuals aged 0–2 yr (n=11), 2–4 yr (n=7), 5–7 yr (n=11), 8–10 yr (n=11), 11–17 yr (n=11), 18–24 yr (n=10), and 25–41 yr (n=10); and U.S. adults (n=8). Statistical analysis: two-way ANOVA with Tukey's multiple comparisons test.

| | *Pf* Healthy baseline | | | HA Healthy baseline | | | *Pf* Convalescence | | | HA Convalescence | | | Diagnostic *Pf* PCR (healthy baseline) | Sex | Age |
|---|---|---|---|---|---|---|---|---|---|---|---|---|---|---|---|
| **Donor ID** | **atBC** | **actBC** | **MBC** | **atBC** | **actBC** | **MBC** | **atBC** | **actBC** | **MBC** | **atBC** | **actBC** | **MBC** | | | |
| ANON3001 | 16 | 6 | 15 | 14 | 8 | 36 | 34 | 15 | 25 | 25 | 29 | 44 | unknown | female | 12 |
| ANON3002 | 1 | 1 | 9 | 20 | 7 | 26 | 3 | 6 | 6 | 14 | 11 | 38 | unknown | male | 12 |
| ANON3003 | 5 | 3 | 9 | 14 | 1 | 5 | 9 | 12 | 5 | 34 | 6 | 4 | negative | female | 12 |
| ANON3004 | 1 | 6 | 9 | 7 | 1 | 28 | 8 | 12 | 8 | 7 | 1 | 17 | negative | male | 13 |
| ANON3005 | 6 | 1 | 17 | 13 | 1 | 6 | 7 | 7 | 15 | 5 | 3 | 6 | negative | female | 13 |
| ANON3006 | 5 | 1 | 8 | 16 | 1 | 6 | 4 | 3 | 2 | 8 | 3 | 1 | negative | female | 11 |
| ANON3007 | 2 | 1 | 7 | 13 | 14 | 47 | 1 | 3 | 5 | 1 | 11 | 2 | unknown | male | 11 |
| ANON3008 | 10 | 1 | 4 | 18 | 11 | 8 | 13 | 11 | 19 | 22 | 11 | 8 | negative | female | 12 |
| ANON3009 | 1 | 1 | 2 | 4 | 1 | 5 | 1 | 2 | 5 | 3 | 2 | 2 | unknown | female | 12 |
| ANON3010 | 2 | 0 | 22 | 12 | 2 | 8 | 7 | 16 | 14 | 24 | 6 | 21 | negative | female | 13 |
| ANON3011 | 0 | 0 | 0 | 0 | 0 | 0 | 6 | 7 | 15 | 10 | 6 | 21 | negative | male | 11 |
| ANON3012 | 0 | 1 | 0 | 3 | 1 | 7 | 2 | 8 | 2 | 3 | 7 | 3 | positive | male | 13 |
| ANON3013 | 0 | 0 | 0 | 0 | 0 | 0 | 2 | 2 | 3 | 2 | 5 | 5 | unknown | male | 12 |
| ANON3014 | 0 | 2 | 6 | 2 | 1 | 2 | 7 | 9 | 8 | 2 | 8 | 6 | negative | male | 12 |
| ANON3015 | 0 | 1 | 2 | 3 | 4 | 11 | 6 | 10 | 10 | 6 | 8 | 10 | negative | male | 13 |
| ANON3016 | 0 | 0 | 0 | 0 | 0 | 0 | 6 | 16 | 21 | 0 | 3 | 16 | positive | female | 14 |
| | total libraries sequenced: 65 | | | | | | total libraries sequenced: 91 | | | | | | | | |

**Table S1.** Cell counts by donor (n=16), timepoint, cell subset, and antigen specificity as sorted for transcriptome analysis of *Pf*- and HA-specific B cell subsets. Samples with a cell count of zero were excluded from library generation (n=24; highlighted in blue). Libraries that failed quality checks were excluded from sequencing (n=8 at baseline, n=4 at convalescence; highlighted in red). *Pf* infection status by PCR for each subject at the baseline timepoint is shown. All subjects were asymptomatic and afebrile at the baseline timepoint.

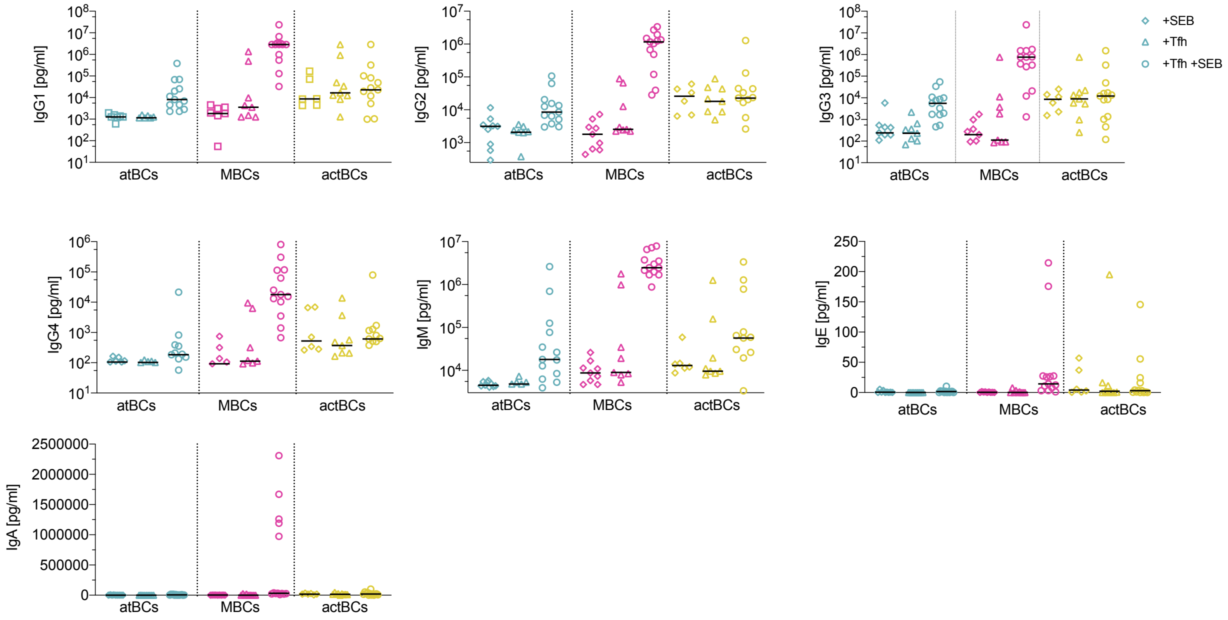


**Supplementary Figure 2: actBCs and MBCs secrete Ig when co-cultured with PD-1⁺CXCR5⁺ CD45RO⁺ CD4⁺ cTfh cells.**

IgG1–4, IgM, IgE and IgA concentration in supernatants after 7-day co-culture with SEB, cTfh cells, or SEB plus cTfh cells. Symbols represent individual donors (n=13), lines represent median. Data are representative of four independent experiments.

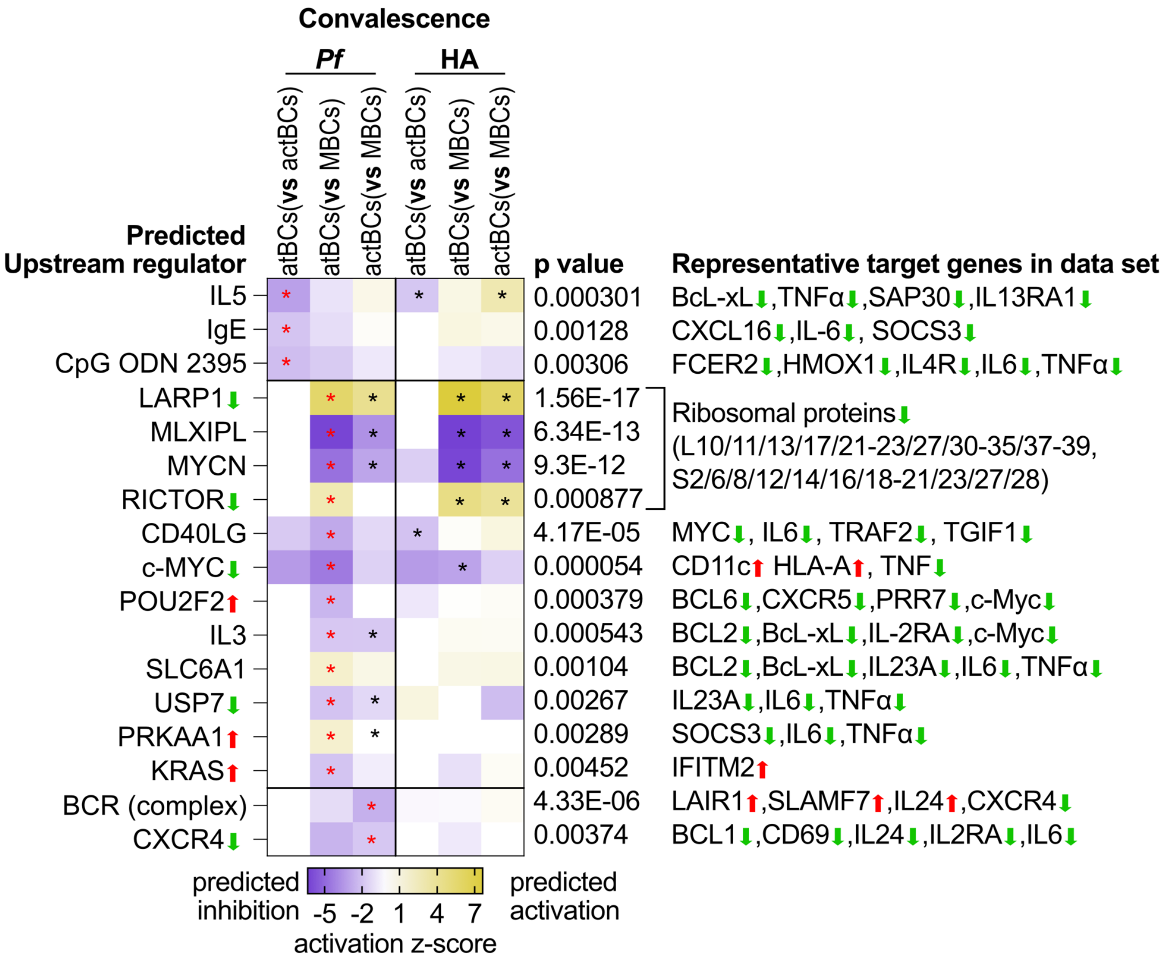


**Supplementary Figure 3: Predicted regulators upstream of transcriptional changes in B cell subsets at convalescence.**

Top upstream regulators identified by Ingenuity analysis using the DEGs of the comparisons atBCs vs actBCs; atBCs vs MBCs; actBCs vs MBCs at convalescence (ΔHB; FDR<0.2, no FC cut-off). Heatmap shows predicted regulators with a |z score|>2 and p<0.005 for the respective comparison with positive z-score indicating activation of the regulator. Listed p values pertain to comparison indicated by red asterisks. Arrow up/down indicates gene expression in the data set.
